# Inhibition of autophagy as a novel therapy for the treatment of neurofibromatosis type 1 tumors

**DOI:** 10.1101/2021.12.20.473481

**Authors:** M. Stevens, Y. Wang, S.J. Bouley, T.R. Mandigo, A. Sharma, S. Sengupta, A. Housden, S. Oltean, N. Perrimon, J.A. Walker, B.E. Housden

## Abstract

Neurofibromatosis type 1 (NF1) is a genetic disorder associated with various symptoms including the formation of benign tumors along nerves. Drug treatments are currently limited. The MEK inhibitor selumetinib is used for a subset of cases but is not always effective and can cause side effects. Therefore, there is a clear need to discover new drugs to target *NF1*-deficient tumor cells. Using a *Drosophila* cell model of NF1, we performed synthetic lethal screens to identify novel drug targets. We identified 54 candidates, which were validated with variable dose analysis as a secondary screen. Five candidates could be targeted using existing drugs, with autophagy inhibitors (chloroquine (CQ) and bafilomycin A1) showing the greatest potential for selectively killing *NF1*-deficient *Drosophila* cells. When further investigating autophagy-related genes, we found that 14 out of 30 genes tested had a synthetic lethal interaction with *NF1*. These 14 genes are involved in the regulation of all aspects of the autophagy pathway and can be targeted with additional autophagy drugs, although none were as effective as CQ. The lethal effect of autophagy inhibitors was conserved in a panel of human *NF1*-deficient Schwann cell lines, highlighting their translational potential. The effect of CQ was also conserved in a *Drosophila NF1 in vivo* model and in a xenografted *NF1*-deficient tumor cell line in mice, with CQ treatment resulting in a more significant reduction in tumor growth than selumetinib treatment. Furthermore, combined treatment with CQ and selumetinib resulted in a further reduction in *NF1*-deficient cell viability. In conclusion, *NF1*-deficient cells are vulnerable to disruption of the autophagy pathway. This pathway represents a promising therapeutic target for *NF1*-associated tumors, and CQ was identified as a promising candidate drug for the treatment of *NF1* tumors.

## INTRODUCTION

Neurofibromatosis type 1 (NF1) is a genetic disorder with autosomal-dominant inheritance affecting 1 in ∼2,700 births (1, 2). Although the penetrance of NF1 is virtually complete after childhood, the disease is characterized by highly variable clinical expressivity. Symptoms include near universal benign, but often disfiguring, peripheral nerve associated tumors known as neurofibromas, as well as malignant tumors, including usually fatal malignant peripheral nerve sheath tumors (MPNSTs) (3). In part reflecting higher rates of vascular defects and cancer, the life expectancy of NF1 patients is reduced by approximately 15 years (4).

NF1 is caused by loss of neurofibromin, a 320 kDa protein whose only widely accepted function is to serve as a RAS GTPase Activating Protein (RASGAP) for H-, K-, N-RAS and R-RAS1, 2, and 3 (5-8). RASGAPs promote the conversion of active RAS-GTP into inactive RAS-GDP by stimulating the low intrinsic rate of RAS-GTP hydrolysis. Consequently, loss of neurofibromin can result in dysregulation of signaling downstream of RAS, the best documented being the RAF/MEK/ERK and PI3K/AKT/mTOR pathways (9). Although dysregulated RAS signaling is believed to be the proximal cause of NF1 symptoms, it is unclear which of the numerous effectors downstream of RAS are relevant for disease progression, as well as the identities of the disease-pertinent targets of the signaling pathways mediating their effects. The situation is further complicated since there is undoubtedly crosstalk between these different pathways. In patients with *NF1*-driven malignant tumors, targeting RAS pathway components such as MEK or ERK is a reasonable therapeutic option, although RAS is subject to highly robust regulation (9), which may explain why, despite considerable effort, effective therapies for RAS-driven cancers have been very challenging to develop. However, chronically blocking RAS may never be an appropriate strategy for treating the many serious but non-life-threatening symptoms of NF1, especially in children.

Currently, there are limited therapies for any *NF1*-associated tumors. The only available drug is the MEK inhibitor selumetinib, which was approved for use in a subset of pediatric plexiform neurofibromas in April 2020. However, not all tumors were responsive to treatment and serious side effects can be associated with MEK inhibition (10-13). Therefore, there is a clear clinical need to discover new drugs that specifically target *NF1*-deficient tumor cells either alone or in combination with selumetinib.

One approach to identify candidate drug targets for tumorigenic diseases is the use of synthetic lethal interaction screens. Synthetic lethal interactions are a type of genetic interaction in which inhibition of either of two genes alone is viable, but the combined inhibition of both genes is inviable. When one of these genes is mutated in tumor cells, such interactions can be exploited to kill those cells exclusively by targeting the synthetic lethal partner gene using a drug (14, 15). This approach is attractive because treatment is expected to be lethal to tumor cells but have no effect on wild-type, healthy cells.

Despite long-term interest in the use of synthetic lethality as a therapeutic strategy to treat tumors, few drugs have successfully progressed to clinical use. A major factor preventing successful development of treatments against synthetic lethal interactions is a lack of consistency between interactions identified in different genetic backgrounds (16). Therefore, candidates identified from a single model system often fail to translate to other model systems and subsequently to clinical use. To overcome this limitation, our approach makes use of diverse model systems by first screening for synthetic lethal interactions with genes mutated in tumors using *Drosophila* cells. The conservation of candidate interactions can then be assessed in a range of other model systems, including human cells, providing a filter to remove interactions that are specific to a single model system. This approach has previously proved successful, leading to the discovery of mizoribine and palbociclib as promising candidates for the treatment of tuberous sclerosis complex (TSC) and Von Hippel-Lindau (VHL)-linked cancers, respectively (17-19). In both cases, hits from *Drosophila* synthetic lethal screens were validated with a high success rate in both human cells and mouse models.

Given the previous success of using the *Drosophila* approach, we have applied this method to identify candidate drug targets to treat *NF1*-deficient tumors. Here, we describe the generation of a *dNF1* null mutant *Drosophila* cell line using CRISPR gene editing and its use in 1) conducting synthetic lethal screens to identify perturbed pathways that confer vulnerability of NF1-deficient cells, and 2) to identify candidate drug targets that might be used for therapeutic benefit to specifically kill NF1-associated tumors. We find dNF1-deficient cells are vulnerable to inhibition of autophagy. Importantly, we show that this selective effect can be reproduced with multiple inhibitors and in several human tumor-derived cell lines, as well as in a *Drosophila in vivo* NF1 model and xenografts of *NF1-*deficient tumor cell lines in mice, indicating that these repurposed drugs may have promise for the treatment of *NF1* tumors. Finally, we show that combined treatment with CQ or bafilomycin A1 and selumetinib results in increased selective killing of *NF1*-deficient cells, indicating the potential for combinatorial therapy.

## RESULTS

### Generation of *Drosophila* and human Schwann cell *NF1* models using CRISPR/Cas9 gene editing

Our previous studies have demonstrated the potential of using cross-species genetic screens to identify candidate therapeutic targets for human disease (17-20). The *NF1* gene is well conserved between *Drosophila* and humans with 68% identity at the amino acid level (**Figure S1**). To use the same approach to find new targets for the treatment of *NF1*-associated tumors, we first used CRISPR gene editing to generate indel mutations in *dNF1* in *Drosophila* S2R+ cells. Sequencing was used to confirm that the induced frame-shift mutations resulted in null *dNF1* alleles (**Figure 1A**). Of note, S2R+ cells are aneuploid (21), and our sequencing results suggest that they have three copies of the *dNF1* gene.

**Figure 1.**
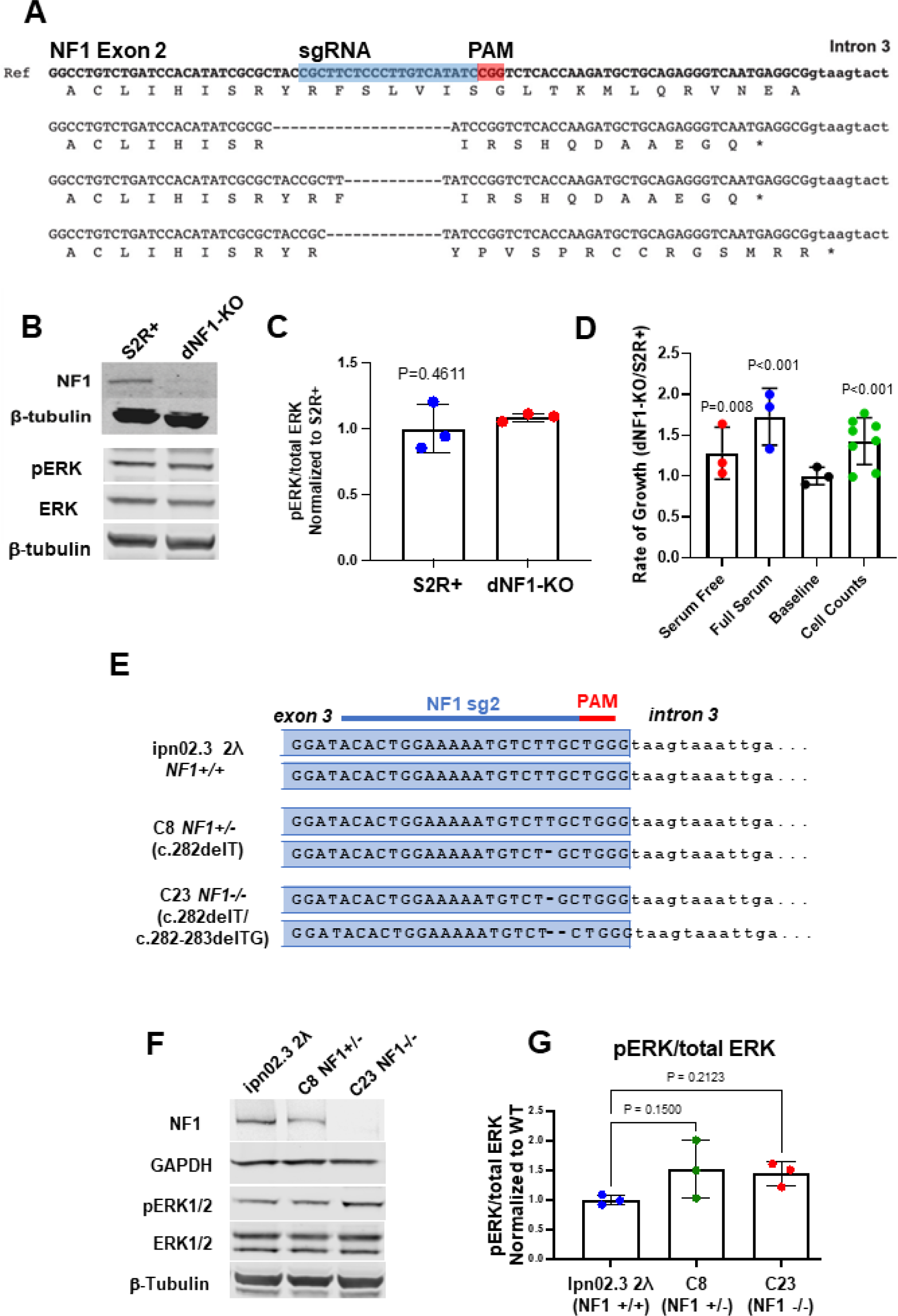
Generation and characterization of NF1-deficient Drosophila and human Schwann cells using CRISPR. **(A)** Drosophila S2R+ cells were transfected with Cas9 and sgRNA designed to target a double-stranded break in exon 2 of dNF1. Molecular analysis of the dNF1-KO line revealed deletions of 20, 11 and 13bp in three alleles (S2R+ cells are aneuploid). The position of the guide RNA is shown in blue and the PAM site in red. Predicted effect of deletions on the amino acids in each allele are shown, each resulting in premature termination. **(B)** Representative image of western blot showing loss of neurofibromin in dNF1-KO cells compared to WT S2R+ cells (two replicates performed) and pERK and total ERK levels in S2R+ and dNF1-KO cells under normal culturing conditions (10% serum). **(C)** Quantification of pERK/ERK ratio in S2R+ and dNF1-KO cells (in 10% serum) shows no significant change (from triplicate experiments). **(D)** Characterization of dNF1-KO cell population growth and proliferation rate as assessed using CellTiter-Glo assays (n=3-8, error bars indicate standard deviation, p values determined using unpaired (except for cell counts, which were paired), two-tailed t-tests) or cell counts. **(E)** CRISPR/Cas9 was used to target exon 3 of NF1 in a telomerase-immortalized human Schwann cell line (hTERT ipn02.3 2λ) (22) to generate isogenic NF1 knock out (NF1-/+ and NF1-/-) cell lines. Line C8 has a heterozygous 1 bp deletion (c.282delT) and line C23 is a transheterozygous combination of 1 bp (c.282delT) and 2 bp deletions (c.282-283 delTG). Position of the guide RNA (NF1-sg2) is shown in blue. **(F)** Western blot analysis of cell lines showing reduction (C8) and absence (C23) of neurofibromin compared to the wild-type ipn02.3 2λ progenitor Schwann cell line. pERK and total ERK levels were assessed in cells grown in 10% serum. **(G)** Quantification of pERK/ERK ratios shows no significant change between wild-type and C8 or C23 (from triplicate experiments).

The resulting dNF1-KO S2R+ cell line (hereafter called dNF1-KO) was characterized by assessing the expression of neurofibromin using western blots. We found no detectable signal in the dNF1-KO line compared to parental wild-type (WT) S2R+ cells (**Figure 1B, Figure S2A**). However, we found no significant increase in pERK levels in dNF1-KO cells under normal culturing conditions, suggesting that other growth conditions (i.e., reduced serum) may be necessary to exacerbate differences in RAS signaling, as is necessary in other NF1 cell line models (**Figure 1B-C, Figure S2B**). Given that neurofibromin is a negative regulator of RAS, we assessed the growth and proliferative phenotypes of dNF1-KO cells compared to S2R+ cells. Consistent with deregulation of a mitogenic pathway, dNF1-KO cells showed an increased rate of growth as measured using CellTiter-Glo assays to assess total ATP levels in the population. This effect was observed in both the presence and absence of serum in the culture media (**Figure 1D**), indicating that culture growth is both accelerated in the absence of dNF1 and is decoupled from upstream growth factor signaling pathways. To determine whether this increase in culture growth was due to increased proliferation, increased cell growth, or both, we performed cell counts following culture in full serum and CellTiter-Glo assays on normalized numbers of cells from each genotype (baseline readings). Cell counts for dNF1-KO showed an increase in cell numbers following culture compared to S2R+ cells, and the ‘baseline’ CellTiter-Glo showed no difference (**Figure 1D**). This suggests that the difference in culture growth is primarily due to increased cell proliferation rather than an increase in the cell size or ATP content of the cells. Together, these results indicate that the dNF1-KO line represents a novel *dNF1* null mutant cell model, with properties consistent with known effects of NF1 loss.

We also utilized CRISPR/Cas9 gene editing to generate indel mutations within exon 3 of *NF1* in wild-type immortalized human Schwann cells (ipn02.3 2λ). Sequencing of single cell clones was used to confirm out-of-frame deletions in one or both *NF1* alleles, resulting in *NF1*-deficient cell lines (**Figure 1E**). The resulting heterozygous (C8) and homozygous (C23) cell lines were characterized by assessing neurofibromin expression using western blots (**Figure 1F, Figure S2C**). Consistent with what we observed in dNF1-KO cells, there was no significant increase in pERK expression under normal culturing conditions (**Figure 1F-G, Figure S2D)**. These results suggest that these otherwise isogenic human cell lines are an appropriate model to validate any results obtained in our dNF1-KO cell lines.

### Mapping synthetic lethal interactions in dNF1-KO cells using a genome-wide RNAi screen

We used a genome-wide dsRNA library to screen in both S2R+ and dNF1-KO cells for synthetic lethal interactions (**Figure S3**). Correlation coefficients ranged between 0.9 and 0.99 (average 0.93) for control wells and between 0.55 and 0.66 (average 0.61) for non-control wells, illustrating a high rate of reproducibility between replicates. Next, we identified synthetic lethal interactions by filtering the results for dsRNA reagents that reduced the viability of dNF1-KO cells (median Z<-1.5) to a greater extent than wild-type cells (median Z⩾-1.5). This analysis identified 134 candidate genes (**Table S1**).

Genetic screens are often associated with false-positive results due to off-target effects from dsRNA reagents or noise in the screen assay. To remove potential false positives, we overlaid the screen hits onto a protein-protein interaction network from the String database (23). Synthetic lethal interactions are generally similar between genes that have related functions; therefore, proteins that physically interact are expected to share synthetic lethal interactions. Using the combination of physical and genetic interaction data, we could remove false positives from the screen results by isolating only hits that have physical interactions with at least one other hit from the genetic interaction screen. In addition, we filtered the candidates to isolate only those with clear orthologs in humans. Following this process, 54 high-confidence candidate targets remained, corresponding to 74 human genes (**Figure 2A, Table S2**).

**Figure 2.**
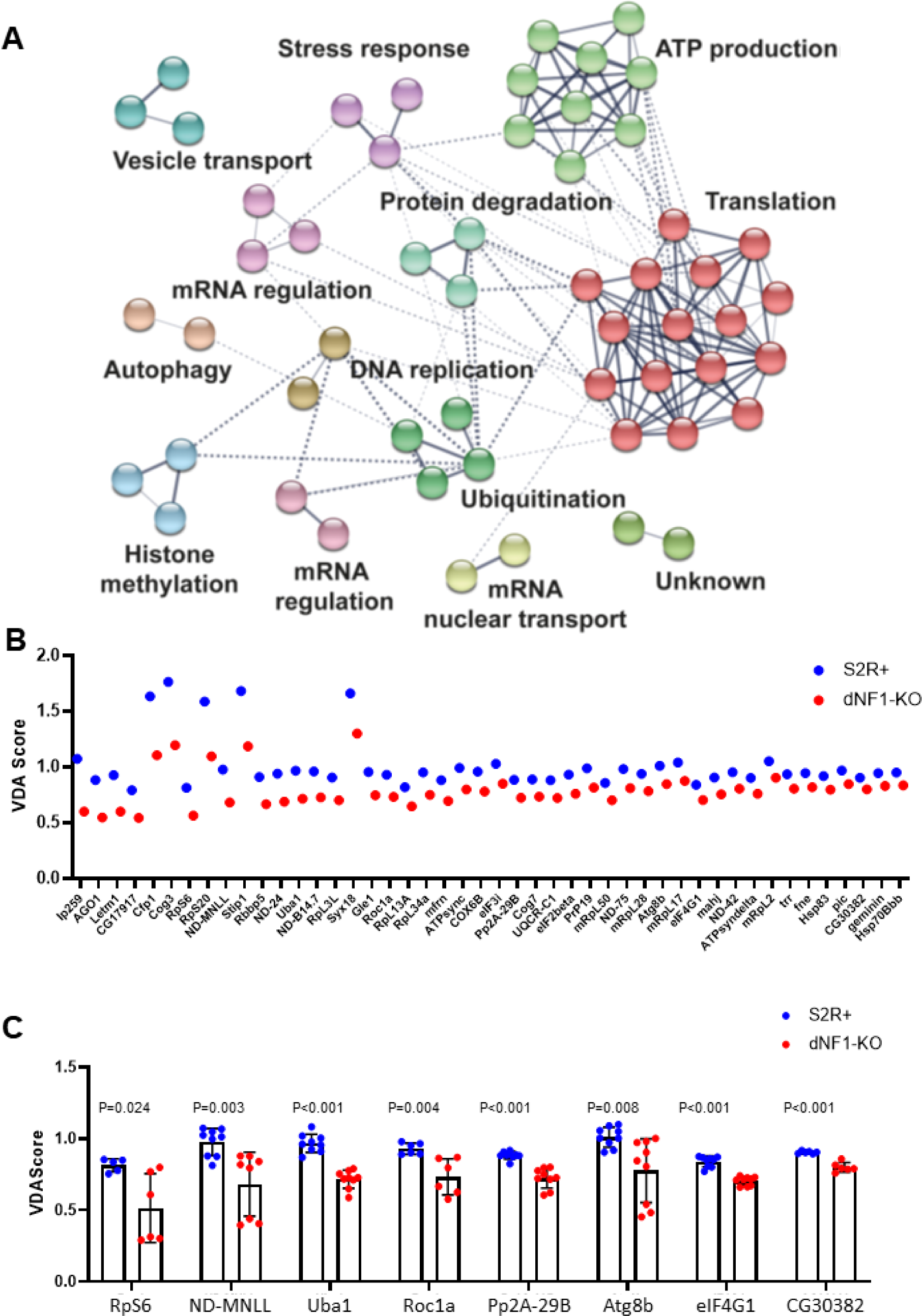
A network of synthetic lethal interaction for Drosophila dNF1 and VDA analysis of candidate drug targets. Using wild-type S2R+ and dNF1-KO cells, we used a near genome-wide dsRNA library to screen approximately 10000 genes for difference in viability when expression is knocked down as assessed by CellTiter-Glo assays, resulting in 134 genes identified as having a synthetic lethal interaction with NF1. **(A)** A synthetic lethal interaction network for dNF1 in Drosophila S2R+ cells generated using Cytoscape (24). Solid lines represent physical interactions within functional groups and dashed lines represent physical interactions between functional groups. **(B)** VDA assays were performed for all 54 candidate genes, with two shRNAs per gene, in WT S2R+ (blue) and dNF1-KO (red) cells. The best shRNA from the 46 genes that reduced dNF1-KO viability by >10% relative to S2R+ controls ranked in order of effect are shown. **(C)** shRNA knockdowns of candidate genes that resulted in a >10% reduction in dNF1-KO viability compared to S2R+ controls and were druggable targets (n=6-9, error bars indicate standard deviation). All eight shRNAs showed a significant reduction in dNF1-KO viability relative to S2R+ controls assessed using two-tailed, unpaired t-tests. Results in panel C are reproduced from panel B for clarity.

### Validation of candidate synthetic lethal interactions using Variable Dose Analysis (VDA)

We used the VDA assay as an additional combinatorial screen to assess synthetic lethality between *dNF1* and all 54 candidate drug targets. Two shRNAs targeting each of the genes were generated. These reagents were tested in S2R+ and dNF1-KO cells. Of the 54 genes, 46 showed a >10% reduction in viability in dNF1-KO cells compared to S2R+ controls (**Figure 2B**; ranked in order of effect on dNF1-KO viability). These results indicate that the network is a reliable representation of the synthetic lethal interaction profile of the *dNF1* gene.

To identify potential drugs for repurposing to treat NF1 tumors, we filtered the candidate gene list to those that could be targeted using existing drugs, resulting in eight candidate drug targets (**Figure 2C**). We then removed candidates that had previously been studied in relation to NF1, leaving five candidate drug targets. We also included MEK (selumetinib) as a control. Note that three drugs that target autophagy were included, and some drugs inhibited multiple targets. In total we tested seven candidate drugs inhibiting six different targets (**Table 1**).

**Table 1.** Candidate genes that selectively decreased dNF1-KO viability that can be targeted with drugs, either directly or through pathway inhibition.

| Drug | Target (direct or indirect): DM (Human) |
| --- | --- |
| Chloroquine* | Autophagy; Atg8b (GABARAP), RpS6 (RPS6) |
| EAD1 | Autophagy; Atg8b (GABARAP), RpS6 (RPS6) |
| Selumetinib* | Dsor1 (MEK, MAPKK) |
| Metformin* | ND-MNLL (NDUFB1) |
| PYR41 | Uba1 (UBA1) |
| LB100 | Pp2A-29B (PPP2R1A and PPP2R1B) |
| Bafilomycin A1 | Autophagy; Atg8b (GABARAP), RpS6 (RPS6) |

\*FDA-approved

### Existing inhibitors selectively affect *dNF1*-KO *Drosophila* cells

Repurposing existing drugs represents the most efficient route to develop new therapeutics. Each of the seven drugs that target the *dNF1* synthetic lethal partner genes were first tested in S2R+ and dNF1-KO cells using CellTiter-Glo assays to measure viability after 48 h of treatment (**Figure S4)**. Of the seven drugs tested in *Drosophila* cells, we selected the two autophagy inhibitors (CQ and bafilomycin A1) for further study, as both showed a significant and consistent effect on reducing dNF1-KO viability.

One of the strongest hits from the genetic screen was *Atg8b*, which encodes a key component of the autophagy pathway. Autophagy is commonly inhibited experimentally using CQ, which is clinically used as an anti-malarial and shows anti-viral properties, and bafilomycin A1. CQ functions to inhibit autophagy by blocking the binding of autophagosomes to lysosomes by diffusing into the lysosomes and altering the acidic environment, thereby inhibiting autophagic lysosomal degradation (25). On the other hand, bafilomycin A1 disrupts autophagic flux by independently inhibiting V-ATPase-dependent acidification and Ca-P60A/SERCA-dependent autophagosome-lysosome fusion (26). We initially focused on CQ because it is generally well-tolerated (27) and can inhibit autophagy *in vivo* at clinically achievable concentrations (28). Bafilomycin A1 is a potent inhibitor of autophagy but is not clinically approved (29). Nevertheless, we tested bafilomycin A1 to provide additional validation of the effects on autophagy inhibition brought about through an independent mechanism.

First, we quantified autophagic flux in dNF1-KO an dS2R+ cells without drug treatment, i.e., how many autophagosomes form and then become degraded, by measuring the difference in the number of autophagic vesicles in the presence versus the absence of a lysosomal inhibitor, CQ. We observed significantly higher levels of autophagic flux in dNF1-KO cells under serum-free conditions compared to S2R+ controls (**Figure 3A**). We then tested whether CQ and bafilomycin A1 would phenocopy the selective effect observed using genetic inhibition of *Atg8b* in *Drosophila* cells. Both S2R+ and dNF1-KO cells were treated with varying doses of CQ or bafilomycin A1 and cell viability was measured using CellTiter-Glo assays, PI staining, and annexin V staining. A significantly greater effect on dNF1-KO cell viability was observed at multiple concentrations of each drug in serum-free media after 48 h of treatment across all three assays, further validating the interaction between autophagy and NF1 and demonstrating that the effect is cytotoxic (**Figure 3B-F**).

**Figure 3.**
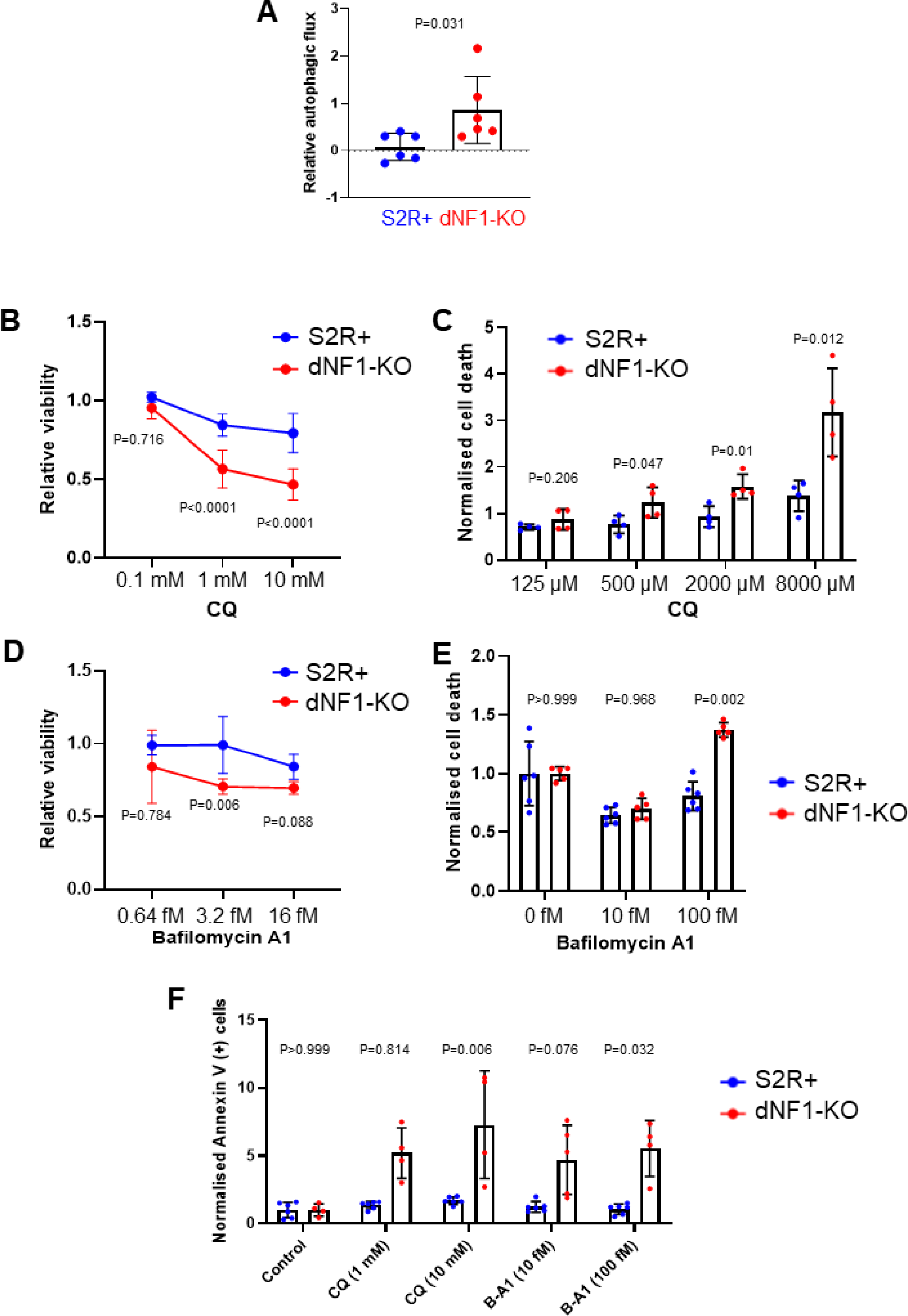
CQ and bafilomycin A1 selectively affect dNF1-deficient Drosophila cells. **(A)** dNF1-KO cells show a significant increase in autophagic flux compared to WT S2R+ control cells in serum-free media as assessed by inhibiting autophagosome flux for 4 h using the lysosomal inhibitor CQ (10 μM), and then measuring the initial rate of accumulation of the fluorescent substrate labelling the autophagosomes using a fluorescent plate reader (normalized to DAPI, and then to S2R+ controls) (n=6, p value obtained using the two-tailed, unpaired, Student’s t-test). **(B)** CQ reduced dNF1-KO cell viability compared to S2R+ cells after 48 h in serum-free media as measured using CellTiter Glo assays (n=4; p values obtained using a two-way ANOVA with Tukey for multiple comparisons). **(C)** Analysis of cell death using PI staining with CQ in S2R+ and dNF1-KO cells grown in serum-free media (n=4; p values obtained using a two-way ANOVA with Tukey for multiple comparisons). **(D)** Bafilomycin A1 reduced dNF1-KO cell viability compared to S2R+ cells after 48 h in serum-free media using the CellTiter Glo assay (n=4; P values obtained using a two-way ANOVA with Tukey for multiple comparisons). **(E)** Analysis of cell death using PI staining with bafilomycin A1 in S2R+ and dNF1-KO cells grown in serum free media (n=6; p values obtained using a two-way ANOVA with Tukey for multiple comparisons). **(F)** CQ and bafilomycin A1 (B-A1) increased annexin V staining in dNF1-KO cells relative to S2R+ controls after 48 h in serum-free media (n=4-6; p values obtained using a two-way ANOVA with Tukey for multiple comparisons). In all cases, bars represent the mean and error bars indicate standard deviation.

### Both early- and late-stage autophagy genes have a synthetic lethal interaction with ***dNF1*, which can be targeted with other autophagy inhibitors**

Autophagy is a complex process involving multiple stages and many different proteins (**Figure 4A**). To determine whether more specific targeting of autophagy components would result in a greater selective effect in *dNF1*-KO cells, we performed an additional VDA screen to assess for synthetic lethal interactions between *dNF1* and 29 key autophagy genes (in addition to *Atg8b*) using S2R+ and dNF1-KO cells (all 29 genes screened are shown in **Table S3**). In total, 14 genes, in addition to the previously identified *Atg8b*, were found to significantly reduce dNF1-KO viability by >10% relative to controls (**Figure 4B**). Interestingly, these 14 genes are implicated across all stages of the autophagy pathway (**Figure 4A**, genes shown in red). Therefore, we tested drugs targeting specific aspects of the autophagy pathway in dNF1-KO cells, including MRT68921 (an early-stage autophagy inhibitor of ULK1 and ULK2), VPS34-IN1 (a potent early-stage autophagy inhibitor of the PI3K-III complex), NSC 185058 (a mid-stage inhibitor of *Atg4b*), CA5f (a late-stage inhibitor of autophagic flux), and obatoclax (a late-stage autophagy inhibitor) (**Figure 4A**). In general, the autophagy inhibitors showed a selective viability effect in dNF1-KO cells, except for MRT68921 and NSC 185058 (**Figure 4C-G**); however, none of the drugs were deemed to be more effective in selectively killing dNF1-KO cells than CQ and bafilomycin A1. CQ also has the advantage of being FDA-approved. Therefore, although targeting autophagy at various stages of the pathway appears to have selective viability effects in dNF1-KO cells, the late-stage autophagy inhibitor CQ could provide the greatest potential for use in the clinic to treat NF1.

**Figure 4.**
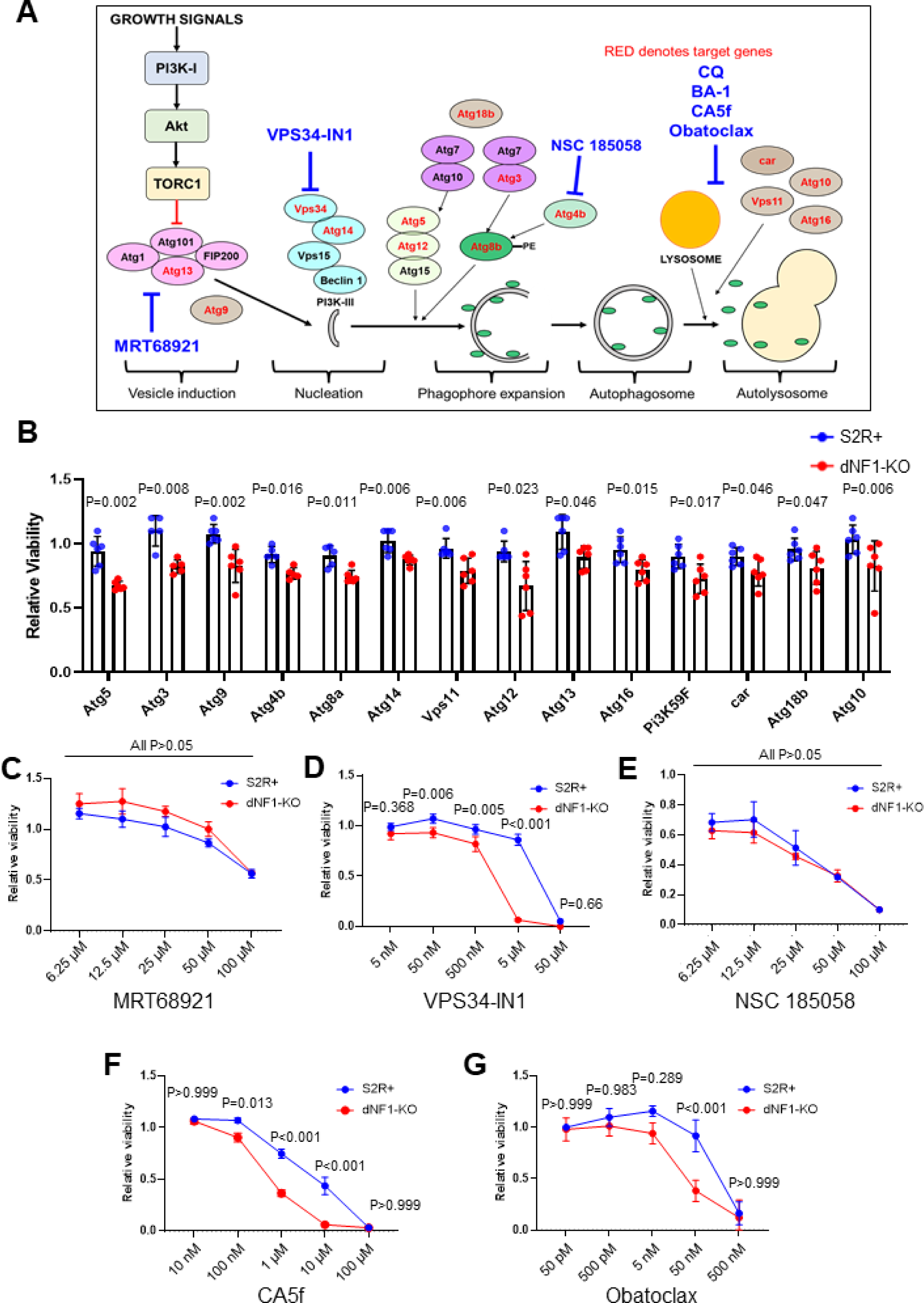
Both early- and late-stage autophagy genes have a synthetic lethal interaction with dNF1, which can be targeted with other autophagy-inhibitors. **(A)** Diagrammatic representation of the autophagy pathway in Drosophila. The genes found to have a synthetic lethal interaction with dNF1 are shown in red. Drugs used to target each stage of the pathway in subsequent viability assays are shown in blue. **(B)** VDA assays performed for 30 autophagy-related genes, with two shRNAs per gene, in S2R+ (blue) and dNF1-KO (red) cells. Shown is the most effective shRNA from each of the 14 genes that reduced dNF1-KO viability by >10% relative to S2R+ controls, ranked in order of effect. **(C-G)** Testing autophagy inhibitors for differences in cell viability between dNF1-KO and S2R+ cells after 48 h in serum-free media as measured using the CellTiter Glo assays (n=4; p values obtained using a two-way ANOVA with Tukey for multiple comparisons). VPS34-IN1 **(D)**, CA5f **(F)**, and obatoclax **(G)** selectively reduced dNF1-KO cell viability compared to S2R+ cells in a dose-dependent manner. By contrast, MRT68921 **(C)** and NSC 185058 **(E)** did not selectively affect dNF1-KO cell viability compared to S2R+ cells. In all cases, data represents the mean and error bars indicate standard deviation.

### CQ and bafilomycin A1 selectively affect *NF1*-deficient human cells

To determine whether the selective effect of CQ and bafilomycin A1 was conserved in human cells, we tested the effects of drug treatment on a panel of human *NF1*-deficient cell lines. These included a pair of otherwise isogenic *NF1^+/-^* (C8) and *NF1^-/-^*(C23) immortalized Schwann cells generated using CRISPR/Cas9 gene editing (**Figure 1F-H**). In addition, we used two pairs of immortalized Schwann cell lines (pair 1: ipnNF95.11C (*NF1^+/-^*) and ipNF95.11b ‘C’ (*NF1^-/-^*) and pair 2: ipnNF09.4 (*NF1^+/-^*) and ipNF05.5 (*NF1^-/-^*)) derived from plexiform neurofibromas (22). Autophagic flux was significantly higher in *NF1^-/-^* cells (ipNF95.11b ‘C’ and ipNF05.5) relative to *NF1^+/-^* controls (ipnNF95.11C and ipnNF09.4), with a similar but not significant effect also observed in the CRISPR-generated *NF1^-/-^*(C23) cells compared to *NF1^+/-^* (C8) controls (**Figure 5A-C**). Similarly, lysosomal activation was increased in ipNF95.11b ‘C’ cells compared to ipnNF95.11C controls, further indicating an increase in baseline autophagy levels (**Figure S5**). Additionally, lysosomal activation was inhibited by CQ and bafilomycin A1 in both *NF1^+/-^* and *NF1^-/-^* cells (**Figure S5**).

**Figure 5.**
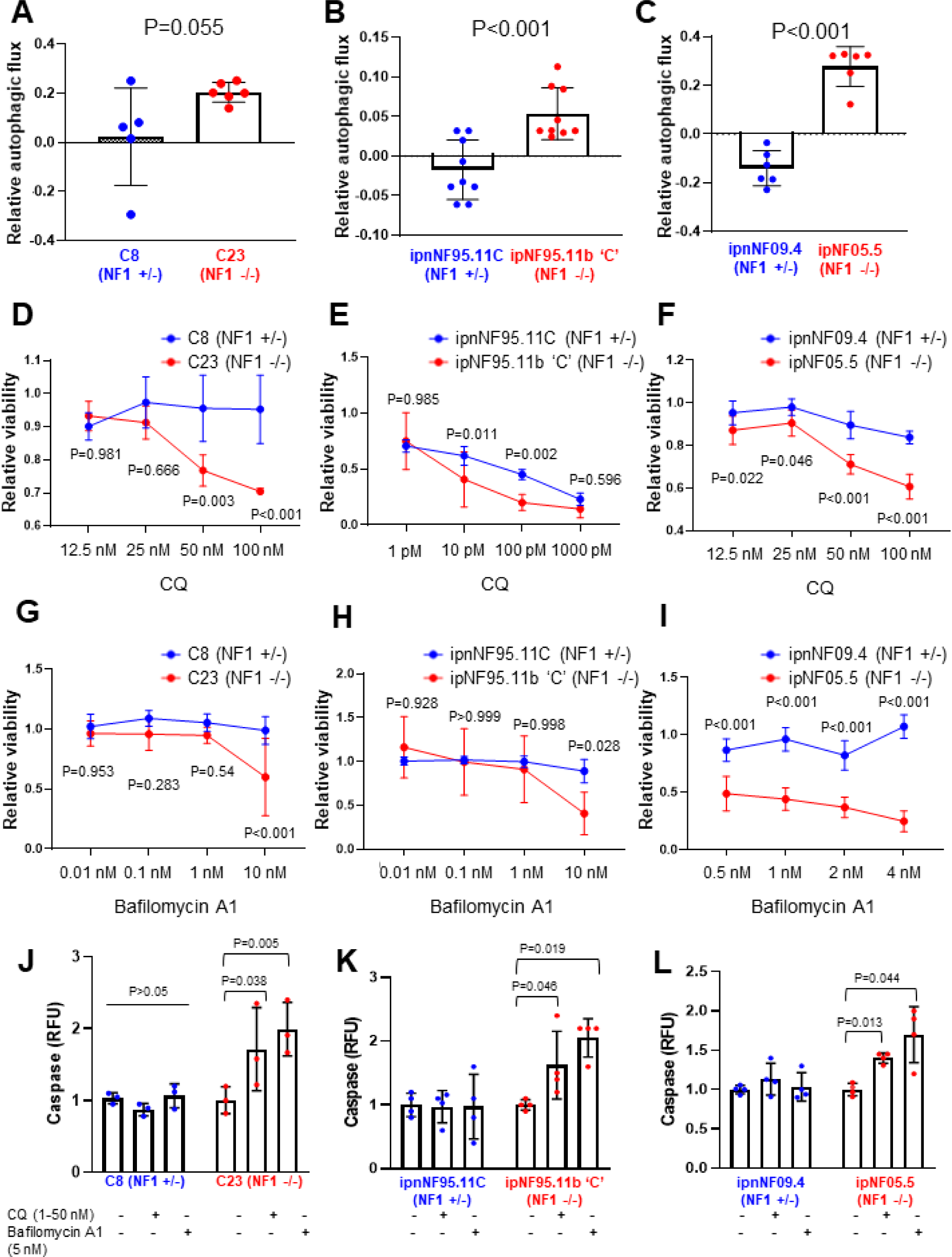
CQ and bafilomycin A1 selectively affect NF1-deficient human cells. (**A-C**) Autophagic flux was increased in C23, ipNF95.11b ‘C’, and ipNF05.5 NF1^-/-^ cells compared to heterozygous controls after 4 h of lysosomal inhibition with CQ (10 μM) in serum-free media (n=5–9; p values obtained using the two-tailed, unpaired, Student’s t-test). CQ significantly reduced NF1^-/-^ cell viability relative to NF1^+/-^ controls at varying doses in C8/C23 cells **(D)**, ipnNF95.11C/ipNF95.11b ‘C’ cells **(E)**, and ipnNF09.4/ipNF05.5 cells **(F)** after 48 h in serum-free media as measured with CellTiter Glo assays (n=3-4; p values obtained using a two-way ANOVA with Tukey for multiple comparisons). **(G-I)** Bafilomycin A1 significantly reduced NF1^-/-^ cell viability relative to NF1^+/-^ controls at varying doses in C8/C23 cells **(G)**, ipnNF95.11C/ipNF95.11b ‘C’ cells **(H)**, and ipnNF09.4/ipNF05.5 cells **(I)** after 48 h in serum-free media as measured with CellTiter Glo assays (n=3-4; p values obtained using a two-way ANOVA with Tukey for multiple comparisons). **(J-L)** We observed an increase in caspase activation in NF1^-/-^ cells relative to NF1^+/-^ control cells when treated with CQ or bafilomycin A1 for 48 h in serum-free media: **(J)**, C8 and C23 cells (50 and 5 nM, respectively), **(K)** ipnNF95.11C/ipNF95.11b ‘C’ cells (1 and 5 nM, respectively), and **(L)** ipnNF09.4/ipNF05.5 cells (50 and 5 nM, respectively) (n=4; p values obtained using a two-way ANOVA with Tukey for multiple comparisons). In all cases, bars represent the mean and error bars indicate standard deviation.

Both CQ and bafilomycin A1 resulted in a significantly greater reduction in the viability of homozygous *NF1^-/-^* deficient cells compared to heterozygous *NF1^+/-^* controls after 48 h of treatment under serum-free media conditions, as measured with the CellTiter-Glo and caspase assays (**Figure 5D-L**), demonstrating that the selective effects are conserved between *Drosophila* and human systems. Although the effective dose of bafilomycin A1 in *Drosophila* cells appears to be very low, human cells were affected by doses in the range of the previously demonstrated IC_50_ of 0.44 nM in human cells (30).

We also tested the additional five autophagy inhibitors in the two patient *NF1*-deficient cell lines (**Figure S6**); however, no drug showed a consistent effect across the panel of human cell lines that was comparable to that of CQ and bafilomycin A1, further highlighting the reproducibility of our *Drosophila* model system.

Together, these results demonstrate that *NF1*-deficient cells have a vulnerability to disruption of the autophagy pathway, which is conserved and reproducible with multiple inhibitors between *Drosophila* and human Schwann cells derived from NF1-associated tumors. Not only does autophagy represent a promising pathway for targeting NF1-associated tumors, but we identified CQ as a candidate drug for the potential treatment of NF1 tumors.

### CQ affects survival in a *Drosophila in vivo NF1* mutant model

As CQ is a well-tolerated FDA-approved therapeutic, and showed a significant effect on dNF1-KO and human NF1^-/-^ cell viability, we chose to take this drug forward to determine synthetic lethality *in vivo*. *Drosophila dNf1* mutant flies show defective Ras signaling, which result in a number of neurobehavioral phenotypes (31-35). For this study, we generated a novel *dNf1* null mutant fly using CRISPR gene editing: *dNf1^C1^* (delAT162-163) (**Figure 6A**). Western blots using lysates prepared from adult heads from *dNf1^C1^* homozygous mutants showed no detectable expression of neurofibromin (**Figure 6B**). In addition, ELISAs showed a 4-fold increase in pERK/ERK of *dNf1^C1^*mutants compared to the WT parental line (**Figure 6C**).

**Figure 6.**
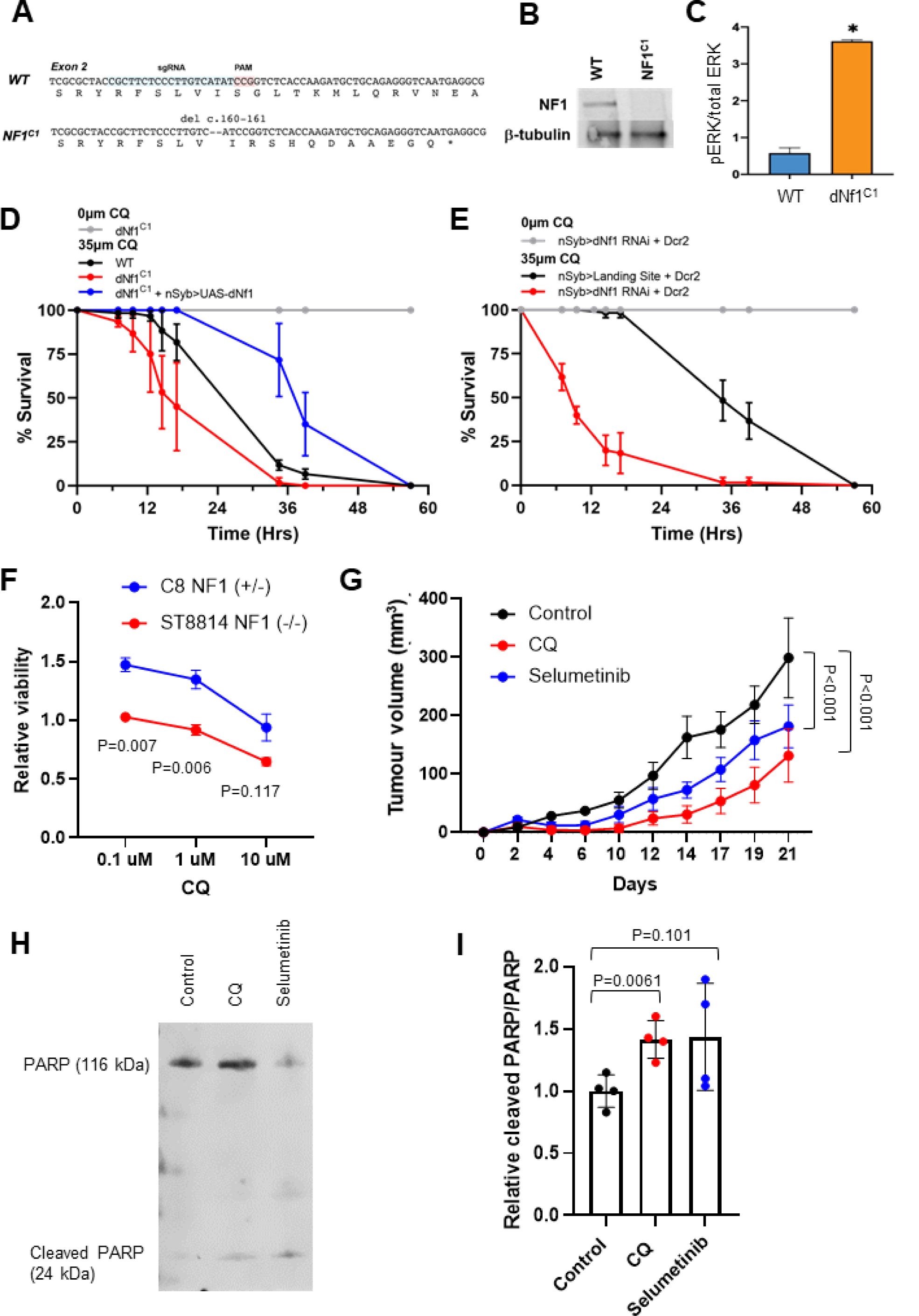
CQ affected lethality in NF1 mutant Drosophila and reduced NF1-deficient MPNST tumor xenograft growth in vivo. **(A)** We generated a novel dNf1 null mutants using CRISPR gene editing: dNf1^C1^ (dNF1 delAT162-163). **(B)** Western blots of anti-dNf1 immunoprecipitates from lysates prepared from adult heads from CRISPR mutants showed no expression of Nf1 in dNf1^C1^ homozygous animals **(C)** ELISA for pERK/ERK showed a 4-fold increase in pERK/ERK of dNf1^C1^ mutants compared to WT flies. Error bars indicate the standard deviation between duplicate samples. A paired, two-tailed t-test was performed to determine significance; *P < 0.05. **(D)** Addition of CQ to 35 µM to food resulted in increased lethality of dNf1^C1^ mutants compared to WT flies (n=3, 20 flies per replicate). The sensitivity of dNf1 mutant flies to CQ was rescued by re-expression of dNf1 from a UAS-dNF1 transgene using the nSyb-Gal4 driver (n=3, 20 flies per replicate) **(E)** dNf1 RNAi knockdown flies (nSyb-Gal4>v109637) show a similarly reduced survival time when cultured on food with 35 µM CQ compared to control flies (nSyb-Gal4>VIE-260B v60100). n=3, 20 flies per replicate. **(F)** CQ significantly reduced NF1^-/-^ mutant cell viability relative to C8 NF1^+/-^ controls at varying doses in ST88-14 cells after 48 h in serum free media, as measured with the CellTiter Glo assay (n=9; P values obtained using a two-way ANOVA with Tukey for multiple comparisons). (**G**) In mice implanted with ST88-14 NF1^-/-^ xenografts, intraperitoneal injections of CQ (50 mg/kg, 3x weekly) or oral gavage with selumetinib (25 mg/kg, 3x weekly) significantly slowed tumor growth compared to vehicle-treated controls. Furthermore, there was a significant reduction in tumor growth in CQ-treated mice compared to selumetinib-treated mice (n=6; P value obtained using a two-way ANOVA). **(H-I)** Following extraction of the xenografts, western blotting revealed the increased protein expression of the 24 kDa fragment of cleaved PARP relative to PARP in CQ-treated mice in comparison to controls (n=4; P values obtained using the unpaired Student’s t-test).

To determine whether CQ affects survival in *NF1*-deficient *Drosophila*, we took two approaches. Firstly, we compared the effect of CQ on *dNf1^C1^* homozygous null mutant flies, the WT parental line, and *dNf1^C1^* with re-expression of dNf1 from a *UAS-dNF1* transgene driven with a pan-neuronal (*nSyb-Gal4*) driver (*dNf1^C1^* + *nSyb-Gal4>UAS-dNf1)*. CQ resulted in increased lethality of *dNf1^C1^* mutants compared to the WT control in a dose-dependent manner (**Figure 6D, Figure S7**). Furthermore, we were able to rescue the CQ sensitivity of *dNf1^C1^* mutant flies by re-expression of dNf1 from a *UAS-dNf1* transgene (**Figure 6D**). Secondly, we tested flies with pan-neuronal RNAi knock down of dNf1 (using *nSyb-Gal4*) compared to a landing site control on food containing CQ (35 mM). Flies with *dNf1* RNAi knockdown, showed significantly reduced survival time on CQ compared to CQ-treated landing site control flies, and untreated flies, phenocopying the effects seen in *dNf1^C1^* mutant flies (**Figure 6E**). Together, these results demonstrate that *dNf1*-deficient flies have vulnerability to disruption of the autophagy pathway, as shown to be conserved and reproducible in *Drosophila* dNF1-KO cells and human Schwann cells derived from NF1-associated tumors. This further highlights the autophagy pathway as a target for the potential treatment of NF1-associated tumors, with CQ as a candidate drug.

### CQ reduced *NF1*-deficient MPNST tumor xenograft growth *in vivo*

In order to test the effects of CQ *in vivo* in a mammalian model, we used the ST88-14 *NF1^-/-^* MPNST xenograft mouse model. Although some NF1 PN tumor cells have been shown to form xenograft tumors, they are very slow growing and require interactions with the tumor microenvironment (36). Therefore, we first assessed whether CQ selectively killed ST88-14 cells *in vitro* using the CellTiter-Glo assay and found CQ to affect cell viability at concentrations slightly higher than those showing an effect in the previously tested *NF1-*deficient Schwann cell lines (**Figure 6F** compared to **Figure 5D-F**). In addition, we found that the ST88-14 cells were more sensitive to CQ than the C8 heterozygous NF1 Schwann cell line (**Figure 6F**).

In mice implanted with ST88-14 *NF1^-/-^* xenografts, treatment with CQ (50 mg/kg in saline, intraperitoneally, three times per week) or selumetinib (25 mg/kg in saline, oral gavage, three times per week) once the tumors had started to grow, resulted in a significant reduction in tumor xenograft growth over a period of 3 weeks in comparison to vehicle-treated controls (**Figure 6G**). Furthermore, there was a significant reduction in tumor xenograft growth in CQ-treated mice in comparison to selumetinib-treated mice, indicating CQ to have a superior effect on *NF1*-deficient tumor xenograft growth. CQ and selumetinib treatment were found to have no toxicity effects in these mice (**Figure S8**), which was also assessed in a prior study on C57BL/6 mice (data not shown).

CQ and selumetinib were found to induce *NF1*-deficient cell apoptosis *in vitro*; therefore, we used western blotting to assess the cleavage of a key apoptosis protein, PARP, in xenografts from control, CQ, and selumetinib-treated mice. We found a significant increase in the 24 kDa fragment of cleaved PARP/PARP in xenografts from CQ-treated mice (**Figure 6H-I**), indicating that CQ was causing cell apoptosis in the NF1 xenografts, thus slowing tumor growth. This increase was also observed in selumetinib-treated mice, although the difference was not significant.

### Selumetinib enhances to viability effect of CQ and bafilomycin A1

Selumetinib, a MEK1/2 inhibitor, is currently the only FDA-approved drug for the treatment of tumors associated with neurofibromatosis type 1 (37). We treated S2R+ and dNF1-KO cells with selumetinib (10 µM) with and without CQ (1 mM) or bafilomycin A1 (100 pM) for 48 h in serum-free media and performed CellTiter-Glo assays to measure cell viability (**Figure 7A**). When selumetinib was combined with CQ or bafilomycin A1, we saw a further reduction in dNF1-KO, but not S2R+ viability relative to the CQ/bafilomycin A1 only, indicating that selumetinib enhances the effects of CQ/bafilomycin A1 on dNF1-KO cell viability.

**Figure 7.**
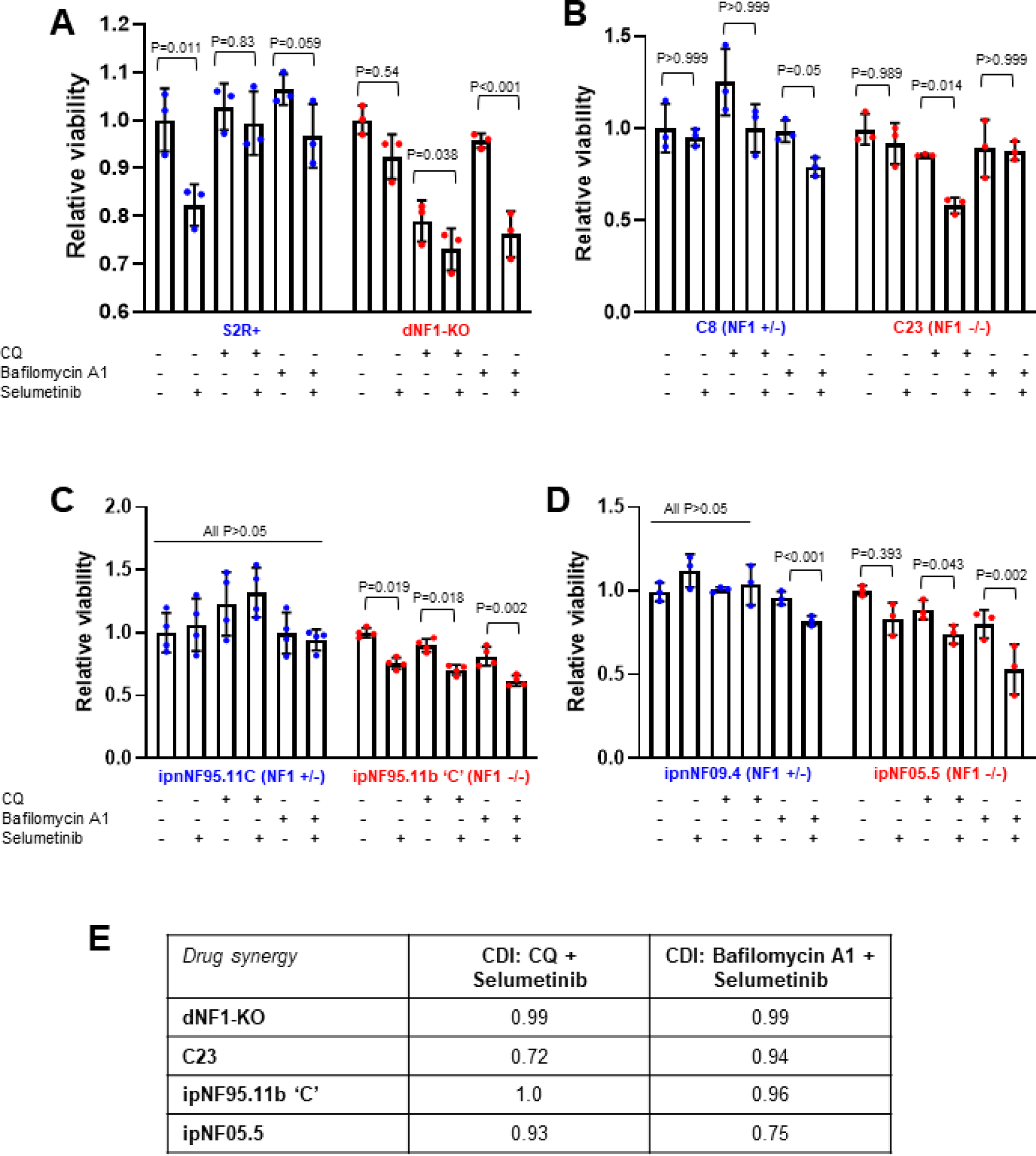
Combined effect of selumetinib with CQ or bafilomycin A1 on NF1-deficient cell viability. **(A)** S2R+ and dNF1-KO cells were treated with 0 µM or 10 µM selumetinib combined with CQ (1 mM) or bafilomycin A1 (10 pM). Alone, selumetinib significantly reduced the viability of S2R+ but not dNF1-KO cells after 48 h in serum-free media, as measured using CellTiter Glo assays. In combination with CQ or bafilomycin A1, selumetinib resulted in a significantly greater reduction in dNF1-KO cell viability compared to CQ alone, which was not observed in S2R+ cells (n=3; p values obtained using a two-way ANOVA with Tukey for multiple comparisons). **(B)** In a similar experiment in human cell lines, selumetinib alone had no effect on the viability of C8, C23 (10 μM), ipnNF95.11C (5 µM), ipnNF09.4, and ipNF05.5 (10 µM) **(B-D)**; however, it did significantly decrease viability of ipNF95.11b ‘C’ NF1^-/-^ (5 µM) **(C)**. Similar to observations in Drosophila cells, the effect of CQ was enhanced when combined with selumetinib in all three NF1^-/-^ cell lines (50 nM, 100 pM, and 50 nM, respectively), whereas the effect of bafilomycin A1 (100 pM, 1 nM, and 1 nM, respectively) was enhanced by selumetinib only in the patient-derived NF1-deficient cell lines (ipNF95.11b ‘C’ and ipNF05.5). (n=3-4; p values obtained using a two-way ANOVA with Tukey for multiple comparisons). In all cases, bars represent the mean and error bars indicate standard deviation. **(E)** Drug synergy between CQ + selumetinib or bafilomycin A1 + selumetinib was calculated using the coefficient of drug interaction (CDI). CDI<1 indicates synergy (with CDI<0.7 indicating a strong synergistic effect), CDI=1 indicates additivity, and CDI>1 indicates antagonism. In this case, drug pairs were classed as synergistic if CDI<0.95.

Similarly, in C8/C23, ipnNF9511.C/ipNF95.11b ‘C’, and ipnNF09.4/ipNF05.5 cell lines, selumetinib significantly enhanced the reduced viability effect of CQ in *NF1^-/-^,* but not *NF1^+/-^*cells (**Figure 7B-D**). Therefore, combined treatment of CQ and selumetinib had a greater impact in *NF1*-deficient cell viability compared to either drug alone. In addition, selumetinib significantly enhanced the viability effect of bafilomycin A1 in patient *NF1^-/-^*(ipNF95.11b ‘C’ and ipNF05.5) but not *NF1^+/-^* cells (ipnNF9511.C and ipnNF09.4), although not in C8/C23 cells (**Figure 7B-D**).

To determine whether these combined effects were additive or synergistic, the coefficient of drug interaction (CDI) was calculated for each cell line treated with CQ + selumetinib or bafilomycin A1 + selumetinib. A CDI<1 indicates synergy, CDI=1 indicates additivity, and CDI>1 indicates antagonism. We observed a relatively strong synergistic effect with CQ + selumetinib in C23 cells, and with bafilomycin A1 + selumetinib in ipNF05.5 cells. Weaker evidence of synergy was also seen in other cell lines (**Figure 7E**).

## DISCUSSION

In this study, we used a *Drosophila* cell culture model of NF1 to identify new potential therapeutic targets and subsequently candidate drugs for the selective killing of *NF1*-deficient cells. Using genetic screening of *Drosophila* cells, we identified 54 candidate synthetic lethal genes that when knocked down resulted in the death of *NF1-*deficient cells without impacting the viability of wild-type, healthy cells. Of these genes, 85% (46/54) were validated in secondary assays, indicating the high quality of the screen results. A key outcome from this screen was the identification of *Atg8b* as a synthetic lethal partner of *NF1*. Atg8b (human ortholog: *GABARAP*) is an autophagy-related protein located in the autophagosome. Autophagy is the process where damaged organelles and unfolded proteins are sequestered into autophagosomes in the cytoplasm, which then undergo fusion with lysosomes (autolysosomes), resulting in degradation of the intracellular components (38). This is important in the regulation of cell growth, maturation, and death.

In cancer, autophagy is reported to take on two opposing roles: 1) degradation of damaged organelles and recycling of macromolecules to maintain a stable cellular environment, which prevents the formation of tumors (39); and 2) aiding in cancer cell survival in response to growth-limiting conditions, contributing to tumorigenesis (40). Furthermore, autophagy has been widely reported to promote cancer cell survival through a drug resistance mechanism (40, 41). Therefore, targeting the autophagy pathway is a potential therapeutic option in the treatment of cancer.

Autophagy is a complex process involving many different genes. In addition to *Atg8b*, *Uba1* was also identified in the screen, which is also linked to autophagy (42). While the selective effects of inhibiting *Atg8b* and *Uba1* were consistent and reproducible in secondary assays, it was surprising that more autophagy linked genes were not identified in our screens. Upon further investigation, we found that approximately half of the 30 autophagy genes tested with low throughput assays showed the selective effects observed with *Atg8b*. These genes have roles from early-stage vesicle induction to late-stage fusion of the autophagosome and lysosome. These results suggest that the entire autophagy pathway is dysregulated in *dNF1*-KO cells, and it is likely that other autophagy genes were missed by the screen due to ineffective RNAi reagents or noise.

Interestingly, *Atg5,* which is involved in the induction of autophagy via autophagic vesicle formation, was found to have the most significant synthetic lethal interaction with *NF1.* Previous studies have implicated ATG5 upregulation in RAS-induced autophagy and malignant cell transformation (43, 44). However, at present, there are no commercially available inhibitors of ATG5. Further studies are required to determine the exact autophagy genes that are dysregulated in our panel of human *NF1*-deficient cell lines.

In total, we assessed seven autophagy inhibitors, and all reproduced the selective effects in *NF1*-deficient cells. However, there was some variation in effects with some drugs not being effective across all cell lines tested. CQ appears to be the most promising of the drugs that we tested for clinical application. It is already approved for clinical use and produced the most consistent and robust effects across the different NF1 cell models that we tested. CQ is a widely used anti-malarial that functions to inhibit autophagy by blocking the binding of autophagosomes to lysosomes by diffusing into the lysosomes and altering the acidic environment, thereby inhibiting autophagic lysosomal degradation (25). Previous studies have shown that CQ has anti-tumor properties in several types of cancer, including glioblastoma (45, 46), hepatocellular carcinoma (47), prostate cancer (48), breast cancer (49), and pancreatic cancer (50). In addition, a recent study reported that the expression of metalloproteinase 1 (MMP1) was down-regulated in *NF1*-deficient fibroblasts, with a further reduction associated with lysosomal degradation of MMP1. Interestingly, treatment of *NF1*-deficient cells with CQ restored MMP1 expression via two mechanisms: activation of the AHR/ERK pathway to enhance the mRNA and protein expression of MMP1, and inhibition of the lysosomal degradation of MMP1 (51). Although this study did not assess whether autophagy was dysregulated in the *NF1*-deficient fibroblasts and did not assess Schwann cells or cells derived from NF1 tumors, it does highlight the therapeutic potential of autophagy inhibition. Similarly, we found that CQ significantly reduced dNF1-KO cell viability relative to wild-type S2R+ cells across a range of concentrations under conditions where autophagy is induced (serum starvation of cells and *NF1* deficiency). This effect of CQ was conserved across a panel of three human *NF1*-deficient cell lines, two of which were derived from neurofibromas from NF1 patients. Furthermore, CQ altered the survival of *dNf1*-deficient *Drosophila in vivo* and reduced the growth of *NF1*-mutant xenografts in mice to an even greater extent than selumetinib. The doses of CQ and selumetinib administered to the mice were deemed non-toxic, as has been widely reported previously (52, 53). Through the inhibition of autophagy with CQ, we observed an increased level of apoptosis in the CQ-treated xenografts, as assessed with the levels of cleaved PARP. Therefore, CQ shows great potential as a therapeutic agent for *NF1*-associated tumors.

While there is a vast array of evidence for the efficacy and safety of CQ, the underlying mechanisms of the tumor suppressive actions of CQ remain to be determined. One potential mechanism is that under starvation conditions, such as those used in this study (i.e. serum starvation), a reduction in glucose transport results in a release of mTOR inhibition of the ULK1 complex, inducing vesicle nucleation and facilitating the process of autophagy. Inhibition of the lysosome using CQ has been shown to inhibit tumor growth and induce tumor cell death *in vitro* (54, 55). We speculate that *NF1*-deficient cells are more susceptible to autophagy inhibition with CQ because the baseline levels of autophagy are higher. RAS has previously been shown to regulate autophagic flux (56). In addition, cancers associated with *RAS* mutations have been reported to be dependent on autophagy, although this appears to be tumor cell line dependent (57, 58). Therefore, cancers with aberrant RAS activity can be more susceptible to autophagy inhibition with CQ. Neurofibromin functions as a RASGAP (GTPase-activating protein), which facilitates RAS inactivation by enabling its GTPase activity (59). In *NF1*-associated tumors, neurofibromin expression is downregulated or absent, resulting in aberrant RAS activity and upregulation of the PI3K/Akt/mTORC1 pathway. Therefore, aberrant RAS activity is expected to negatively regulate autophagy; however, RAS is implicated in many signaling pathways and so it has a multifaceted role in autophagy regulation. For example, RAS also activates the RAF1/MEK1/2/ERK and RAC1/MKK7/JNK signaling pathways, both of which are known to activate autophagy (56). Furthermore, upregulation of ATG5 and ATG7 has been implicated in RAS-induced autophagy and malignant cell transformation (43, 44). Therefore, we speculate that aberrant RAS activity in *NF1*-deficient cells results in the initiation of autophagy, and that inhibition of this pathway has anti-tumor effects. When combining our data on the effects of CQ and bafilomycin A1, we provide strong evidence for inhibition of the autophagy pathway as a target in the treatment of *NF1*-associated tumors.

Finally, we found that inhibition of autophagy with CQ and bafilomycin A1 increased the sensitivity of *NF1*-deficient cells (both dNF1-KO and human *NF1^-/-^* cells) to MEK1/2 inhibition with selumetinib. Selumetinib is currently the only FDA-approved drug for the treatment of tumors associated with neurofibromatosis type 1 (37). The phase 2 trial (SPRINT) for the use of selumetinib in plexiform neurofibromas reported clinically meaningful improvements in 71% of patients (13), prompting its FDA approval for patients ages 2 to 18 years with NF1 who have symptomatic, inoperable plexiform neurofibromas. However, there are toxicity effects related to long-term MEK inhibition. Interestingly, selumetinib had no selective effect in the dNF1-KO cells compared to wild-type S2R+ controls when used alone. Furthermore, we observed no difference in the phosphorylation of ERK between S2R+ and dNF1-KO cells under baseline conditions, which suggests that RAS activity levels are similar in the two cell lines. Therefore, while there is extensive evidence that RAS activation leads to dependence on autophagy, it is possible that there is a second, RAS-independent, mechanism that also contributes to the selective effect of autophagy inhibition. Such a mechanism may explain why we see a combinatorial effect of autophagy inhibition and MEK1/2 inhibition. This finding is in line with a previous study, which showed that combined inhibition of MEK1/2 plus autophagy had a synergistic anti-proliferative effect in pancreatic ductal adenocarcinoma cell lines, which display aberrant K-RAS activity, as well as patient-derived pancreatic ductal adenocarcinoma xenograft tumors in mice (60).

In conclusion, we have shown using multiple techniques, reagents, and models that inhibition of autophagy has potential as a novel therapeutic strategy for the treatment of NF1 tumors. Given the existing clinical use of CQ and the robust and conserved effects that we observe between cell culture models, this candidate drug has a high chance of successful translation to the clinic, resulting in a positive impact on NF1 patients.

## METHODS

### Cell culture

*Drosophila* Schneider (S2R+) cells, both WT and dNF1-KO, were cultured at 25°C in Schneider’s media (Gibco) containing 1% antibiotic (Gibco) and 10% fetal bovine serum (Gibco). Human cell lines used include: *NF1^+/-^* and *NF1^-/-^* immortalized human Schwann cell (SC) lines, derived from the ipn02.3 2λ cell line using CRISPR/Cas9 gene editing as described below; two pairs of immortalized human SC lines derived from plexiform neurofibromas from NF1 patients (22), which included ipnNF95.11C (*NF^+/-^*) and ipNF95.11b ‘C’ (*NF1^-/-^*) cells (germline *NF1* mutation: c.1756delACTA), and ipnNF09.4 (*NF1^+/-^*) and ipNF05.5 (*NF1^-/-^*) cells (germline *NF1* mutation: c.3456_3457insA). All human cell lines were cultured at 37℃ in 5% CO_2_ in DMEM media (Merck) containing 1% antibiotic (Gibco) and 10% fetal bovine serum (Gibco). For ipnNF09.4 and ipNF05.5 cells, the media was also supplemented with 50 ng/ml neuregulin-1 (NRG-1) (Sigma).

### Generation of dNF1-KO *Drosophila* cells

The dNF1-KO cell line was generated using methods described previously (19, 61). WT S2R+ cells were co-transfected with the pl018 plasmid to express Cas9 and the sgRNA (CGCTTCTCCCTTGTCATATC) and pAct-GFP plasmid to mark transfected cells. Five days after transfection, individual cells with the highest 15% GFP signal were isolated and seeded into wells of 96-well plates using FACS. Single cells were cultured using conditioned media for approximately 3 weeks. DNA was then extracted from each candidate cell population and assessed using HRMA to identify those carrying mutations at the sgRNA target site. Positive candidates were then sequenced to confirm frame-shift mutations were present in each *NF1* allele (**Figure 1A**).

### Generation of *NF1*-deficient human Schwann cell lines using CRISPR-Cas9

The wild-type human hTERT-immortalized Schwann cell line (hTERT ipn02.3 2λ) was a gift from Dr. Peggy Wallace, University of Florida (22). CRISPR-Cas9 genome editing was performed using NF1-sg2 (ACACTGGAAAAATGTCTTGC), a sgRNA targeting human *NF1* exon 3 in the lentiCRISPR v2 plasmid, which was a gift from Dr. Feng Zhang (Broad Institute) (62). hTERT ipn02.3 2λ cells were transfected by electroporation using the Amaxa Basic Nucleofector Kit for Primary Mammalian Fibroblasts (Lonza) and the program U-023, according to the manufacturer’s instructions. After 48 h of selection on puromycin (1 μg/ml) in DMEM/10% FBS, cells were re-plated in to allow single clone isolation. Genotyping of selected single clones was performed using PCR using primers NF1-exon3-FOR (CCCCAATTCAAGATTCTGGT) and NF1-exon3-REV (ATCGCACTCTCCCACAACTC). PCR products were treated with ExoSAP (Affymetrix) and sequenced using NF1-exon3-SEQ (TGCCATTTCTGTTTGCCTTA).

### Synthetic lethal screen

Screens were performed using wild-type S2R+ or dNF1-KO cells using the genome-wide RNAi library from the Sheffield RNAi Screening Facility. In total, 10,000 cells were seeded into each library well in 10 µl serum-free Schneider’s *Drosophila* media. Plates were then incubated at room temperature for 45 minutes before addition of 35 µl media with 10% FBS. Libraries were incubated at 25°C for 5 days before CellTiter-Glo assays (Promega) were performed. Screens were performed in triplicate in each cell line (**Figure S3**).

Data were analyzed by first normalizing all values to the median of each row and column of the library plate to allow direct comparison between plates. Z-scores were then calculated for each RNAi reagent using the average and standard deviation of each replicate screen and correlation between replicates was used to assess the quality of screen results. Reagents were considered hits if the Z-score in at least 2/3 of replicates was below -1.5 in dNF1-KO cells and above -1.5 in S2R+ cells. We note that some assay plates were affected by position effects; these were identified manually and removed from the analysis prior to correlation analysis.

Functional gene groups were determined using the in-build clustering tool in the STRING database to group genes. Functions of each group were determined manually by searching for associated GO terms or assessing annotated functions of human orthologs in cases where the *Drosophila* gene was not sufficiently annotated.

### Variable dose analysis (VDA)

VDA is an RNAi-based method in which each cell within a population receives a different dose of shRNA (20, 63). The relative knockdown efficiency of each cell is then measured with a fluorescent reporter. On the day of transfection S2R+ and dNF1-KO cells were plated at 1 x 10^4^ cell/100 µl culture media, per well of a 96-well plate. Cells were incubated at 25℃ for 40 mins to allow adhesion. Cells were then transfected with 40 ng actin-GFP and 160 ng shRNA expression plasmid using 0.6 µl FuGENE^®^ HD transfection reagent in a total volume of 10 µl. We used a positive (*thread*, an apoptosis inhibitor that induces cell death when inhibited) and negative (*white*, known to have no viability effect in these cells) shRNA on each plate for normalization of data. Plates were then sealed and incubated for 4 days at 25℃ in a humidifying chamber.

Flow cytometry was used to identify GFP-positive cells (transfection efficiency). Area under an inverted cumulative distribution curve was used as a readout of relative viability, normalized to the positive and negative control. More detailed protocol and data analysis information can be found in Sierzputowska et al. (63).

### CellTiter-Glo assays

The CellTiter-Glo assay (Promega; G7570) is a luminescent viability assay based on the quantification of ATP, which is an indicator of metabolically active cells. All cells were plated in white 384-well plates at a seeding density of 5 x 10^3^ cells/25 µl culture media in either complete or serum-free media. S2R+ cells were left to adhere for 40 mins at 25℃ and human cells were left to adhere for 4 h at 37℃ in 5% CO_2_ before treatment. Cells were treated with 250 nl per well of each drug (CQ, bafilomycin A1, and selumetinib) in PBS at varying concentrations in replicates of 5-8 using the Mosquito LV Genomics (SPT Labtech) and incubated for 48 h. The CellTiter-Glo assay was then performed according to the manufacturer’s instructions, and luminescence was measured using a plate reader (TECAN Infinite M200 Pro).

### Autophagy assay and Caspase assay

An autophagy assay kit (Abcam; ab139484) was used to measure autophagic vacuoles in live cells using a dye that selectively labels autophagic vacuoles. All cells were plated in 96-well plates in either complete or serum-free media. S2R+ cells were left to adhere for 40 mins at 25℃ and human cells were left to adhere for 4 h at 37℃ in 5% CO_2_ before treatment. Cells were treated with 10 µM per well of CQ in PBS at varying concentrations and incubated for 4 h. The autophagy assay was then performed according to the manufacturer’s instructions, and fluorescence intensity was measured using a plate reader (TECAN Infinite M200 Pro). Autophagic flux was quantified by inhibiting lysosomal fusion with CQ and then measuring the initial rate of accumulation of autophagosomes using a fluorescent plate reader.

Caspase activity was assessed using the same plating and treatment method, except that cells were treated for 48 h, via the generic caspase activity assay kit (Abcam; ab112130) per the manufacturer’s instructions. In both the autophagy and caspase assay, autophagy/caspase fluorescence was normalized to DAPI in each well.

### Annexin V and PI staining

The Annexin V-FITC apoptosis kit (Abcam; ab14085) was used to detect apoptosis by staining of phosphatidylserine molecules that have translocated to the outside of the cell membrane. Cells were co-stained with propidium iodide (PI) to detect dead cells in the population. S2R+ cells were left to adhere for 40 mins at 25℃ before treatment. Cells were treated with 1 µl per well of each drug (CQ, bafilomycin A1) in PBS at varying concentrations, and incubated for 48 h. The annexin-V/PI assay was then performed according to the manufacturer’s instructions, and fluorescence intensity was measured using flow cytometry.

### Western blotting

Lysates from S2R+ cells, dNF-KO cells, human immortalized Schwann cells, and homogenized xenograft tissue were prepared in RIPA buffer supplemented with protease inhibitors (Complete Protease Inhibitor Cocktail, Sigma) and phosphatase inhibitors (NaF/Na_3_VO_4_/β-glycerophosphate). Lysates for testing expression of neurofibromin were resolved on NuPAGE 3–8% Tris-Acetate and for assessing levels of pERK/ERK on NuPAGE 4-12% Bis-Tris protein gels. After transfer to nitrocellulose membranes, blots were processed according to the Odyssey CLx protocol (LI-COR). Antibodies used for immunoblotting: anti-*Drosophila* neurofibromin (mouse, ascites purified mAb21 and mAb30, 1:500 each), anti-human neurofibromin (mouse, Infixion r07E, 1:1000), anti-phospho-ERK (mouse, Sigma M8159, 1:2500), total ERK (rabbit, CST 9102, 1:1000), anti-β-tubulin (mouse, Developmental Studies Hybridoma Bank E7, 1:10,000), GAPDH (rabbit, Proteintech 10494-1-AP, 1:15,000), and anti-PARP (rabbit, Cell Signaling 952, 1:1000). Secondary antibodies used were anti-mouse Alexa Fluor Plus 800 (goat, Invitrogen, A32730, 1:10,000) and anti-rabbit Alexa Fluor Plus 680 (goat, Invitrogen, A21109, 1:10,000).

### *Drosophila* husbandry and stocks

Flies were cultured on cornmeal/agar food medium according to standard protocol and housed at 25°C.

The *Nf1^C1^* mutant fly line was generated by CRISPR/Cas9 gene editing (64). using the sgRNA line: *y[1] sc[*] v[1] sev[21]; P{y[+t7.7] v[+t1.8]=TKO.GS01796}attP40* (GS01796 sgRNA sequence: CGCTTCTCCCTTGTCATATC) and a germline source of Cas9: *y[1] M{w[+mC]=nos-Cas9.P}ZH-2A w[*]* (Bloomington Stock #54591). The *UAS-dNf1* transgene encodes full-length *dNf1* cDNA corresponding to the RF isoform with addition of introns 9 and 10 and was previously published (65). The RNAi and landing site control lines were obtained from the Vienna *Drosophila* RNAi Center: *dNf1 RNAi* line: P{KK101909}VIE-260B (VDRC #109637) and VIE-260B (VDRC #60100). *UAS-Dicer2* was included to potentiate the RNAi effect (66): *P{w[+mC]=UAS-Dcr-2.D}10* (Bloomington: 24651). Male flies were used for all experiments.

### Assessing drug sensitivity in flies

Flies were raised on standard fly food and kept at 25°C. For testing drugs, adult flies were transferred to Formula 4-24 Instant *Drosophila* Medium (Carolina Biological) prepared using water (control) or chloroquine (chloroquine diphosphate; Sigma C6628) in water. Concentrations of CQ tested ranged from 0-40 µM (Figure S7) and 35 µM was used for the experiments in Figure 6 D and E. For each genotype, three replicates of 20 flies were added to each condition and kept at 25°C. Flies were monitored periodically to assess lethality. Survival was reported as the percent of flies still alive at each time point.

### NF1 tumor xenografts

All experiments were conducted in accordance with UK legislation and with local ethics committee approval (University of Exeter AWERB). Two million ST88-14 cells mixed 1:1 with Matrigel (ThermoFisher Scientific) were injected subcutaneously into the right flank of male CD-1 nude mice (Charles River Laboratories). Tumors were measured three times weekly with a caliper, and the tumor volume was calculated as follows: [(length + width) / 2] x length x width. Once the tumor diameter began to increase over two separate measurements, mice were intraperitoneally injected with either saline or CQ (50 mg/kg in saline) or received selumetinib (25 mg/kg in saline) via oral gavage three times per week (n=6 mice per group). Mice were culled by cervical dislocation (Schedule 1) when the control tumor sizes reached the allowed endpoint (12 mm in diameter), and tumors were dissected. Tumors were flash frozen for further analysis.

## Competing Interest Statement

B.E.H. is a shareholder and founding director of Quest Genetics Ltd. The remaining authors declare no competing interests.

## Acknowledgements

This study was funded by a Medical Research Council project grant (MR/V009583/1) and a Medical Research Council Confidence in Concept (CiC) award (MC_PC_19037) to B.H. and NIH R21 NS096402, DOD-NFRP W81XWH-16-1-0220 and the Neurofibromatosis Therapeutic Acceleration Program (NTAP) (J.A.W.). We would also like to thank Nerve Tumours UK for their support as collaborators on the CiC award. S.J.B was supported by the Dorothy and Spiro Latsis Fellowship for NF1 Research. N.P. is an investigator of HHMI. We thank the Sheffield RNAi Screening Facility, Biomedical Sciences, University of Sheffield, supported by the Wellcome Trust (grant reference number 084757) for providing RNAi libraries.

## Notes

### Summary of Updates

We have added in vivo validation of CQ as a potential treatment for NF1-deficient tumors.

