## Supplementary Figures for "Inhibition of autophagy as a novel therapy for the treatment of neurofibromatosis type 1 tumors"

### Figure S1. The *NF1* gene is well conserved between *Drosophila* and humans with 68% identity at the amino acid level.

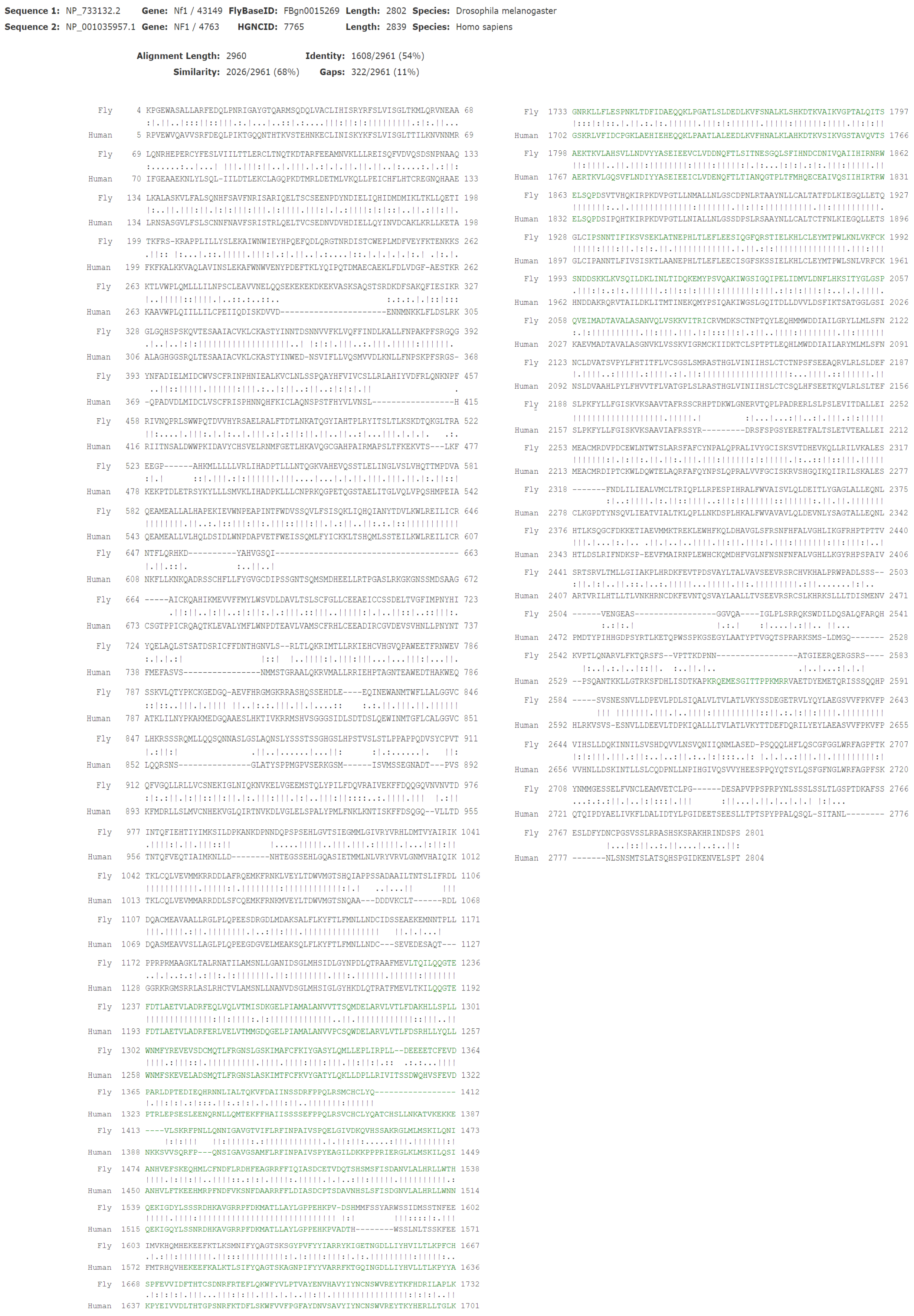

**Figure S2. Western blot data**
**(A)** Relates to Figure 1B. **(B)** Relates to figure 1C-D. **(C-D)** Relate to Figure 1F-G. In each case, a representative image is shown from at least three replicates.

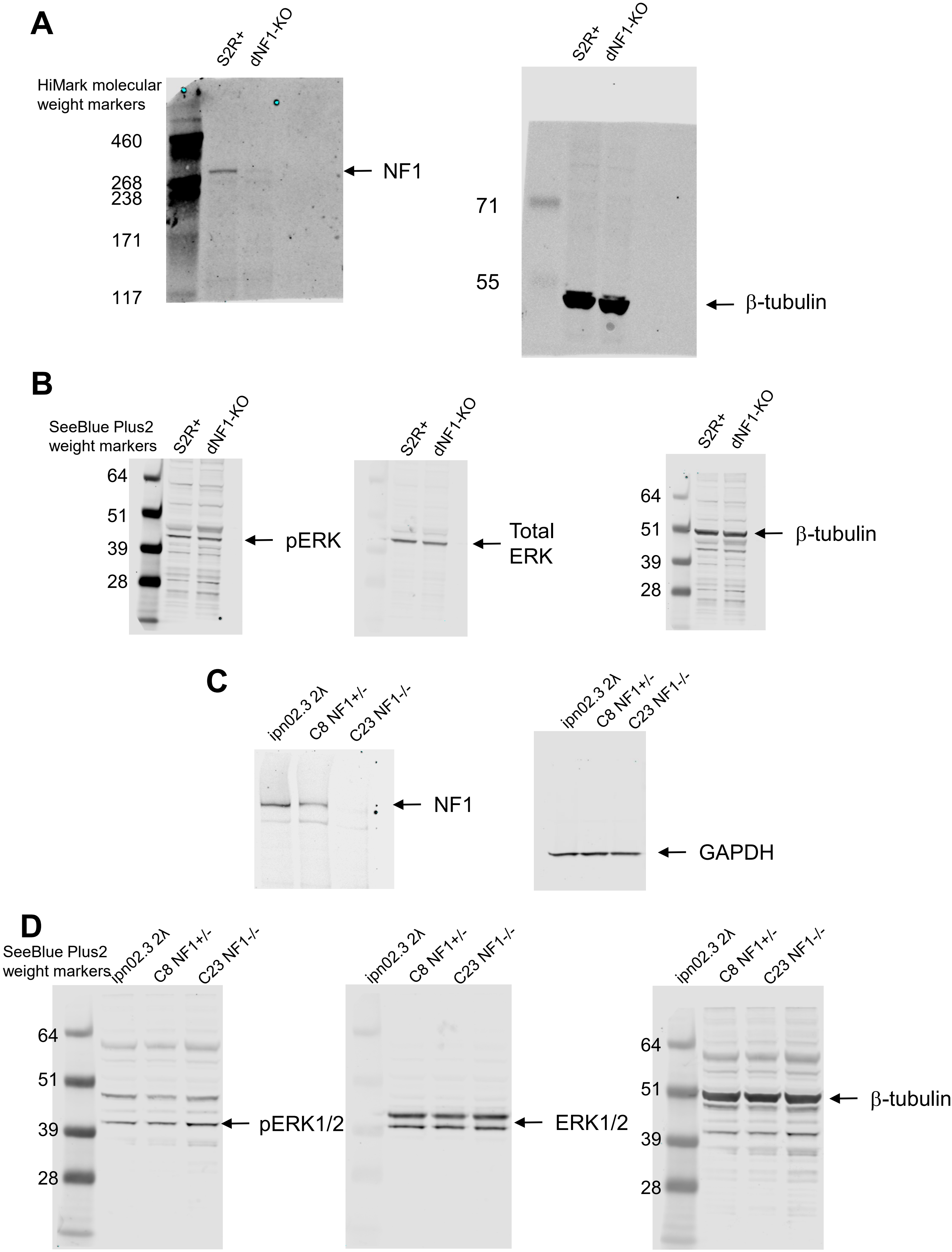

**Figure S3. Identification of synthetic lethal interactions with *NF1.***

**(A)** Using dNF1-KO cells, we used a genome-wide dsRNA library to screen approximately 10000 genes, resulting in 134 genes identified as having a synthetic lethal interaction with *NF1*. **(B)** Summary of results from the synthetic lethal screen. Z-scores are plotted for NF1-KO cells on the vertical axis and WT S2R+ cells on the horizontal axis. The red box indicates the position of reagents selected as hits. **(C)** Correlation analysis of screen replicates indicating high data quality. Each bar represents the correlation coefficient from comparison between two screen replicates in WT or dNF1-KO cells as indicated or between the median Z-scores for each cell line. Bars indicate the mean from each pairwise combination of replicates and error bars indicate standard error of the mean.

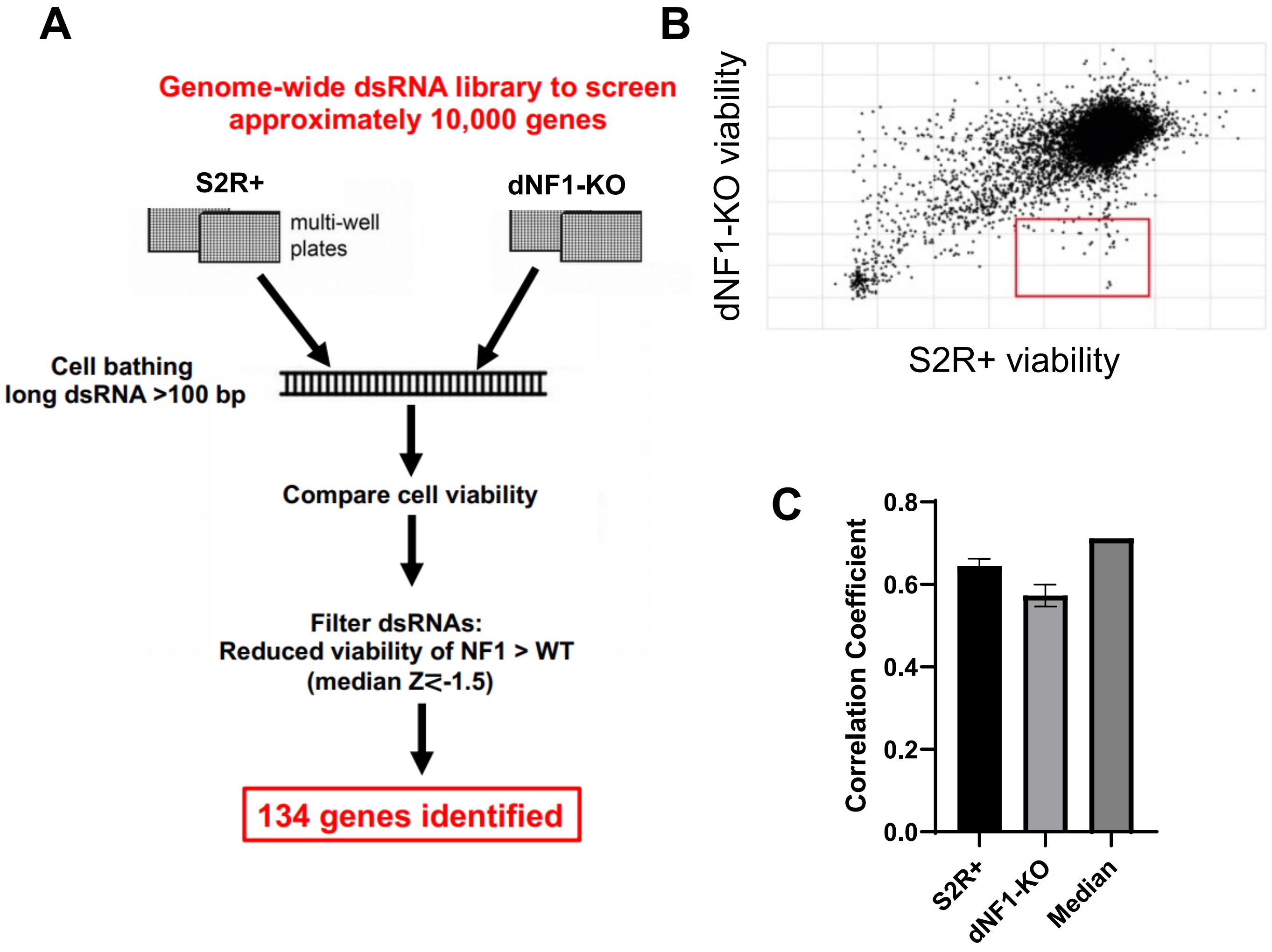

**Figure S4. Drug dose curves. (A)** Dose curves for the seven drugs with *Drosophila* orthologs as targets in S2R+ and dNF1-KO cells. **(B)** Dose curves for CQ and Bafilomycin A1 in human CRISPR/Cas9 genome edited Schwann cells ($NF1^{+/-}$ and $NF1^{-/-}$). **(C)** Dose curves for CQ and Bafilomycin A1 in human patient cells: ipnNF95.11C ($NF1^{+/-}$) and ipnNF95.11b 'C' ($NF1^{-/-}$).

**A**

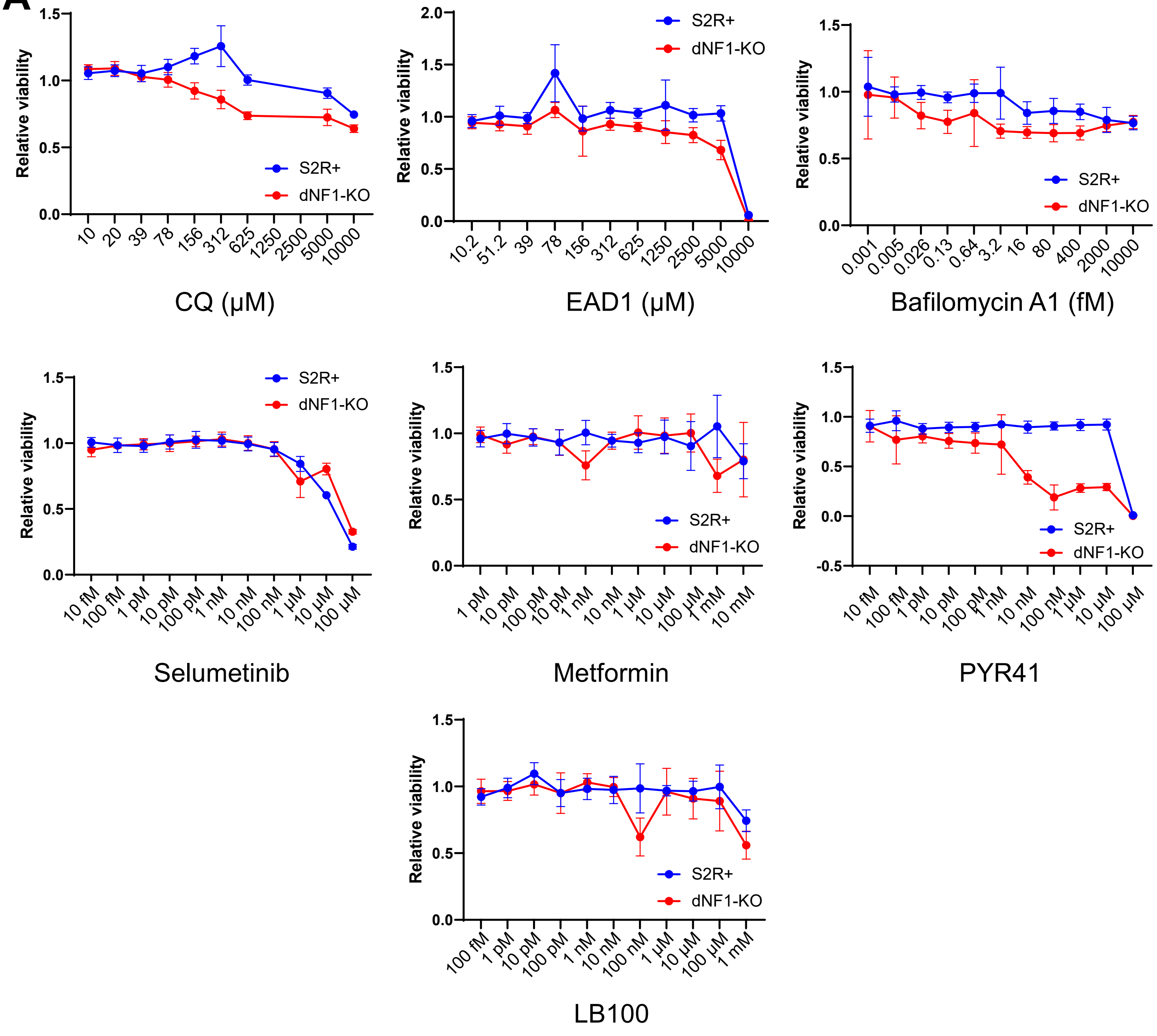

B

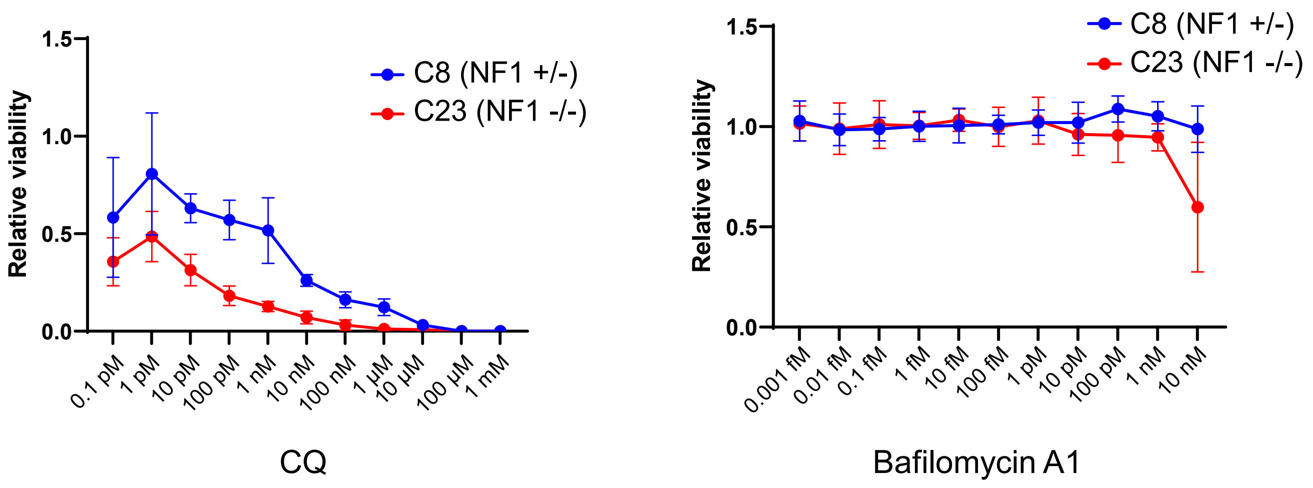
C8 (NF1 +/-)
C23 (NF1 -/-)
Relative viability
1.5
1.0
0.5
0.0
0.1 pM
1 pM
10 pM
100 pM
1 nM
10 nM
100 nM
1 µM
10 µM
100 µM
1 mM
CQ
0.001 fM
0.01 fM
0.1 fM
1 fM
10 fM
100 fM
1 pM
10 pM
100 pM
1 nM
10 nM
Bafilomycin A1

C

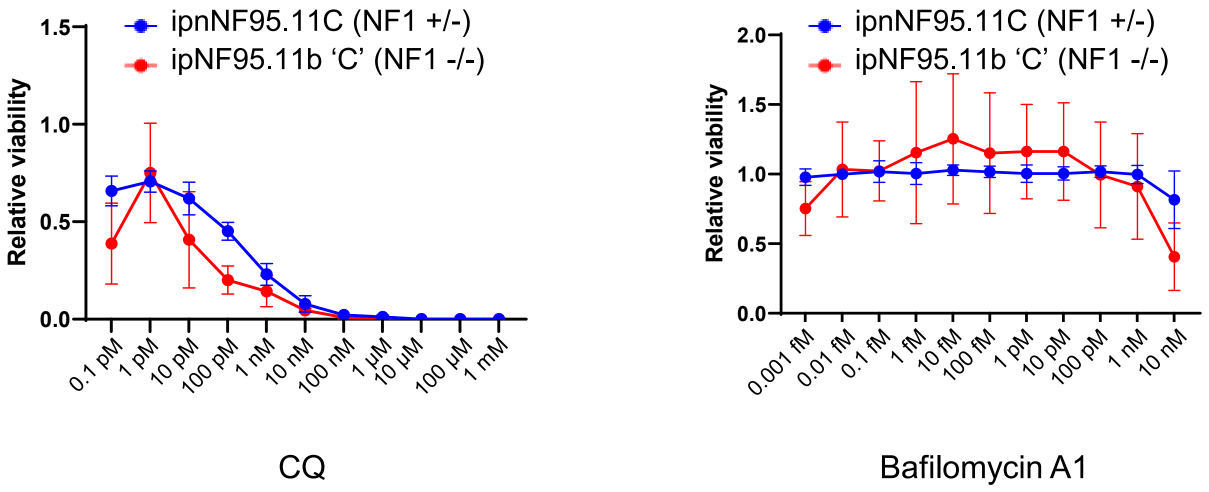
ipnNF95.11C (NF1 +/-)
ipNF95.11b 'C' (NF1 -/-)
Relative viability
2.0
1.5
1.0
0.5
0.0
0.1 pM
1 pM
10 pM
100 pM
1 nM
10 nM
100 nM
1 µM
10 µM
100 µM
1 mM
CQ
0.001 fM
0.01 fM
0.1 fM
1 fM
10 fM
100 fM
1 pM
10 pM
100 pM
1 nM
10 nM
Bafilomycin A1

**Figure S5. Live lysosome activity assay.** ipnNF9511.C and ipNF95.11b 'C' cells were treated with/without CQ or bafilomycin A1 (B-A1) for 2 h in serum free media before measuring lysosomal activity using a lysosome-specific self-quenching substrate (Abcam). The substrate acts as endocytic cargo, and upon degradation, a fluorescent signal is generated. Therefore, fluorescence is proportional to lysosome activity. Increases fluorescence (green) was observed in ipNF95.11b 'C' (NF1 -/-) cells compared to heterozygous controls. CQ and B-A1 inhibited lysosomal activity in both cells. Nuclei are shown in blue (marked with DAPI).

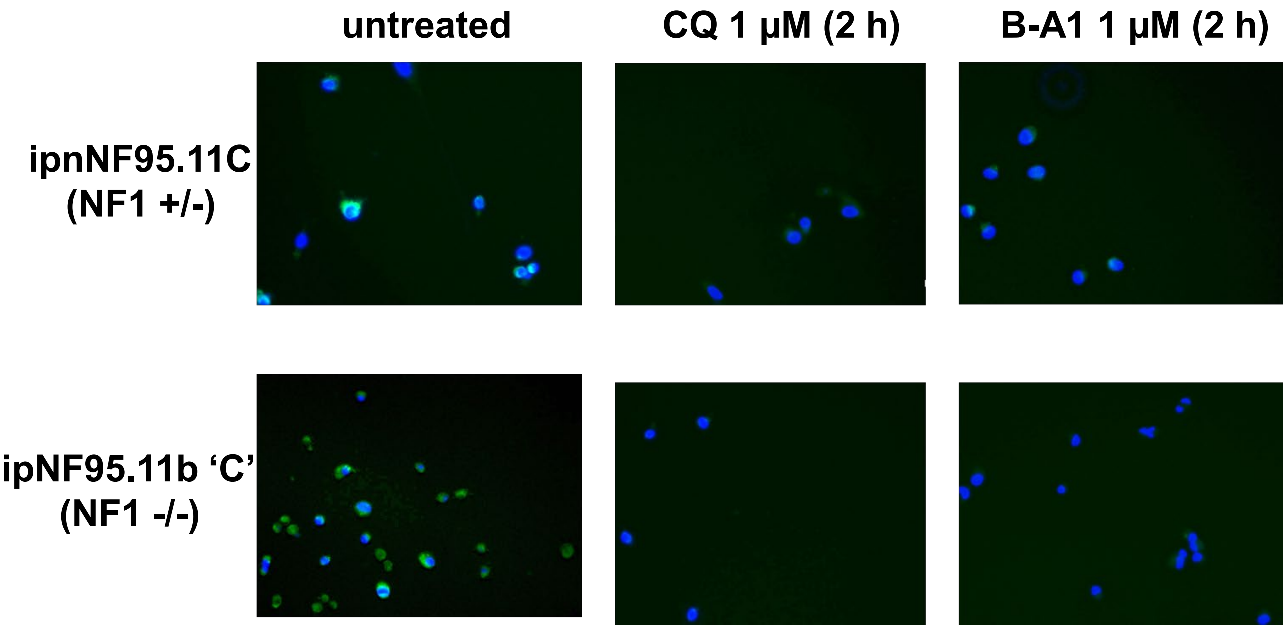

**Figure S6. Drug dose curves. (A)** Dose curves for the five autophagy drugs in patient cells: ipnNF95.11C ($NF1^{+/-}$) and ipnNF95.11b 'C' ($NF1^{-/-}$), and ipnNF09.4 ($NF1^{+/-}$) and ipNF05.5 ($NF1^{-/-}$).

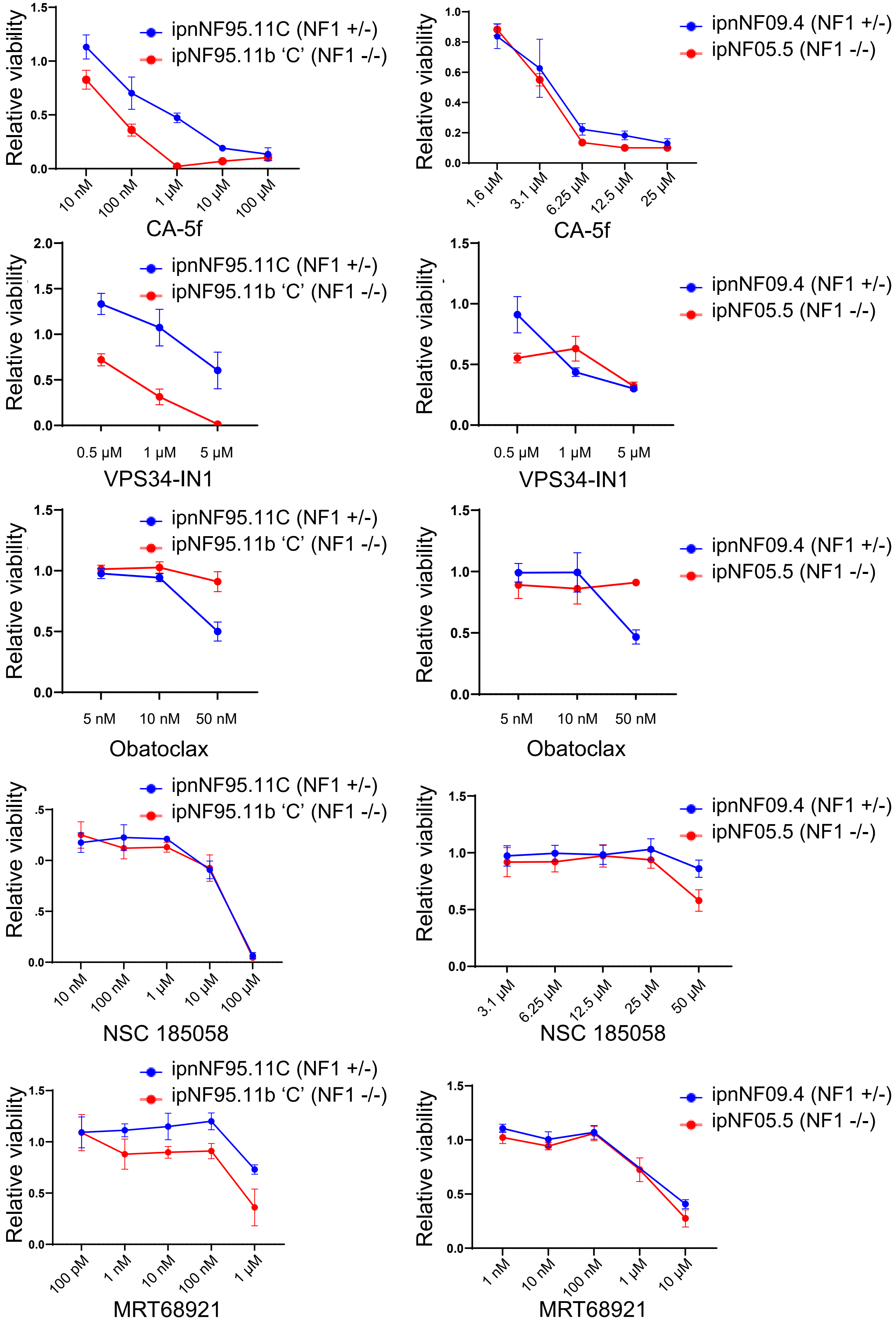

**Figure S7. CQ results in an increased dose-dependent lethality of *dNf1$^{C1}$* mutants compared to *WT* control flies.** Fly food was supplemented with the indicated concentrations of CQ and survival was determined by counting living flies at each time point (n=3, 20 flies per experiment).

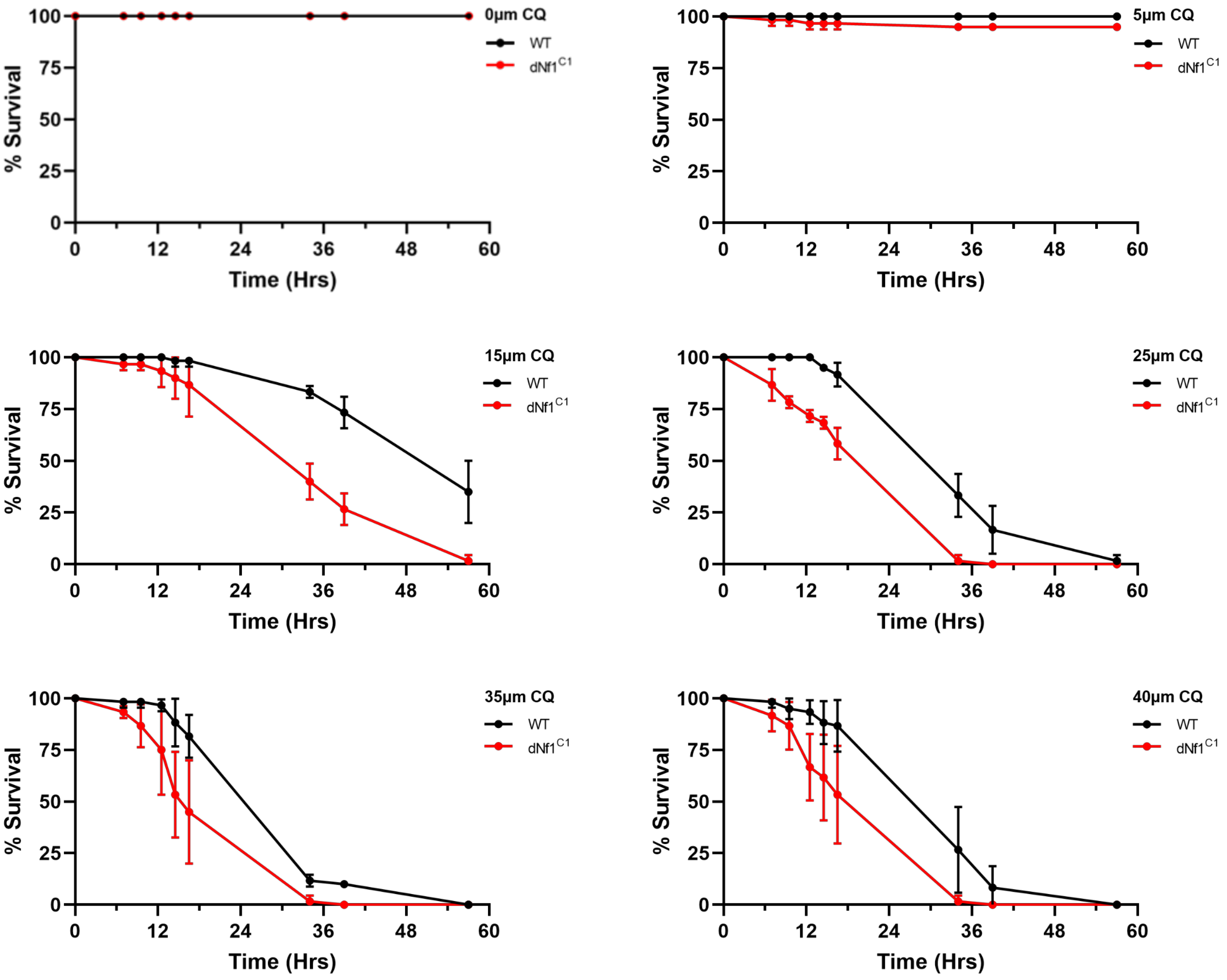

**Figure S8. Mouse weight change during treatment.** The adult mice were weighted three times per week during treatment with either saline (control), CQ, or selumetinib. No mice lost weight during the treatment period, indicating no toxic effects associated with the dose used of CQ (50 mg/kg) and selumetinib (25 mg/kg).

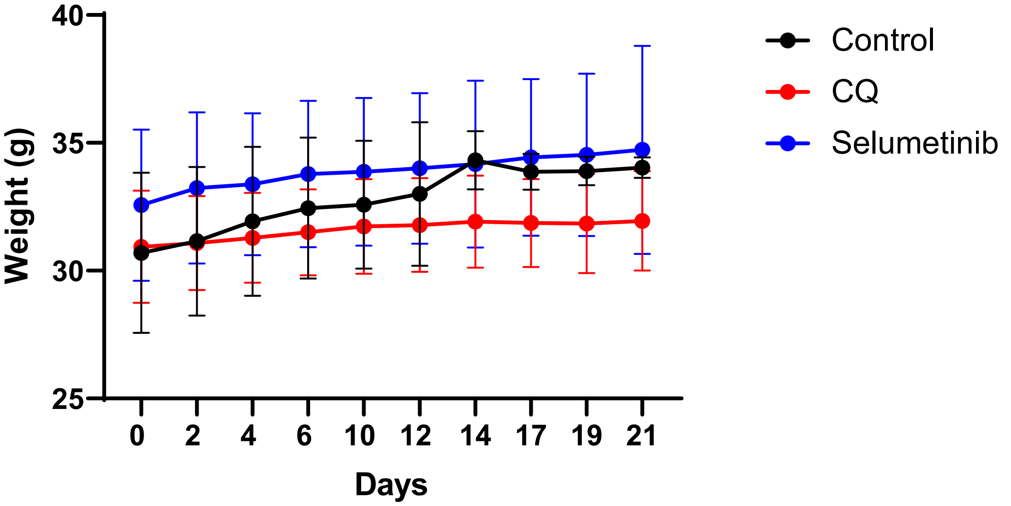
